# Investigating Developmental Changes in Scalp-Cortex Correspondence Using Diffuse Optical Tomography Sensitivity in Infancy

**DOI:** 10.1101/2020.08.22.262477

**Authors:** Xiaoxue Fu, John E. Richards

## Abstract

**Significance:** Diffuse optical tomography (DOT) uses near-infrared light spectroscopy (NIRS) to measure changes in cerebral hemoglobin concentration. Anatomical interpretations of NIRS data requires accurate descriptions of the cranio-cerebral relations and DOT sensitivity to the underlying cortical structures. Such information is limited for pediatric populations because they undergo rapid head and brain development.

**Aim:** The present study aimed to investigate age-related differences in scalp-to-cortex distance and mapping between scalp locations and cortical regions of interest (ROIs) among infants (2 weeks to 24 months with narrow age bins), children (4 and 12 years) and adults (20 to 24 years).

**Approach:** We used spatial scalp projection and photon propagation simulation methods with age-matched realistic head models based on MRIs.

**Results:** There were age-group differences in the scalp-to-cortex distances in infancy. The developmental increase was magnified in children and adults. There were systematic age-related differences in the probabilistic mappings between scalp locations and cortical ROIs.

**Conclusions:** Our findings have important implications in the design of sensor placement and making anatomical interpretations in NIRS and fNIRS research. Age-appropriate realistic head models should be used to provide anatomical guidance for standalone DOT data in infants.

## Introduction

Diffuse optical tomography (DOT) uses near-infrared light spectroscopy (NIRS) to measure changes in cerebral hemoglobin concentration (Huppert, Franceschini, & Boas, 2009). DOT does not provide anatomical information about the location of the hemodynamic signal. Spatial scalp projection can be implemented to interrogate the brain region(s) underlying the scalp sensor (i.e. optodes) locations (Tsuzuki & Dan, 2014). An alternative approach is to generate a forward model of DOT sensitivity using photon propagation simulations (Boas, Culver, Stott, & Dunn, 2002). The forward model can then guide DOT image reconstruction to recover the brain locations of hemoglobin concentration changes (Cooper et al., 2012; Culver, Siegel, Stott, & Boas, 2003). There are considerable brain structural changes during infancy through childhood and adulthood (Richards & Conte, 2020; Richards & Xie, 2015). Accurate models of DOT sensitivity must account for age-related changes in the head (Brigadoi, Aljabar, Kuklisova-Murgasova, Arridge, & Cooper, 2014). The present study examined age-related changes in scalp-to-brain distance and the correspondence between scalp locations and anatomical regions. We used age-appropriate, realistic head models with extensive coverage of the infancy period (2 weeks through 2 years), two child ages (4, 12 years), and young adults (20 to 24 years).

### Scalp-to-Cortex Distances Measured by Scalp Projection Methods

The initial step to understanding the underlying cortical structures being measured with NIRS is to measure the distance from the NIRS recording locations to the cortical surface. The 10-20, 10-10, and 10-5 systems provide a standardized and reproducible method for placing electrodes based on cranial landmarks and are often used to identify NIRS optode recording positions (Jurcak, Tsuzuki, & Dan, 2007). Spatial scalp projection uses an algorithm to project an electrode location on the scalp down to a location on the cortical surface (Okamoto & Dan, 2005; Tsuzuki & Dan, 2014). Adult studies projected 10-20 (Okamoto et al., 2004) and 10-10 electrodes (Koessler et al., 2009) to standard brain templates. Okamoto et al. (2004) found that the scalp-to-cortex distance was shallower in the frontal, temporal and occipital regions, but deeper in the parietal and along the intra-hemispherical fissure. Spatial scalp projection facilitates the understanding of the sampling depth required to measure the cortical activities using NIRS recording or DOT.

It is important to examine age-related changes in scalp-to-cortex distance. There is extensive brain morphological development from infancy to adolescence (Gilmore et al., 2011; Makropoulos et al., 2016; Richards & Conte, 2020; Richards & Xie, 2015). One change is the distance from the scalp to cortical landmarks (Heschl’s gyrus, inferior frontal gyrus, frontal pole, occipital pole, parieto-occipital sulcus, and vertex). There are significant increases in scalp-to-cortex distances from newborn to age 12 (Beauchamp et al., 2011). Kabdebon et al. (2014) virtually placed 10-20 electrodes on individual MRIs from 3- to 17-week-old infants. The distance between the scalp electrode positions and the cortical surface decreased from the frontal to occipital locations. The pattern was not observed in adult head models (Okamoto et al., 2004). Studies adopting the scalp projection method indicate that the scalp-to-cortex distance is smaller in infants (Emberson, Crosswhite, Richards, & Aslin, 2017; Kabdebon et al., 2014) and children (Whiteman, Santosa, Chen, Perlman, & Huppert, 2017) than in adults (Okamoto et al., 2004). Existing infant studies sampled a single age, or a narrow age range. Hence, there is insufficient information about the differences across age on overall scalp-to-cortex distance or scalp-to-cortex distance by electrode positions.

### Scalp-location-to ROI Mapping Using Scalp Projection Methods

The spatial scalp projection method can also be used to identify the correspondence between scalp electrode locations and the underlying anatomical regions of interest (ROIs). The electrode-location-to-ROI mapping can be established first by transforming the individual’s own MRI and electrode locations to a canonical average MRI template which has an anatomical stereotaxic atlas. Scalp electrode locations can then be projected to the cortical surface to identify to the corresponding ROI(s). Adult studies have demonstrated that there was a relatively consistent correspondence between the 10-20 (Okamoto et al., 2004) and 10-10 (Koessler et al., 2009) electrode positions and the underlying macro-anatomical ROIs. A methodological challenge in the ROI mapping is the limited availability of subjects’ own MRIs. Co-registration methods for standalone DOT data have been developed so that electrode positions on the subject’s scalp can be transformed to the space of reference MRIs in a database (Fekete, Rubin, Carlson, & Mujica-Parodi, 2011; Tsuzuki et al., 2012). The co-registration methods result in similar results as using subjects’ own MRIs (Tsuzuki et al., 2012; Tsuzuki & Dan, 2014).

Co-registering scalp electrode locations with reference MRIs is an effective method for identifying electrode-location-to-ROI correspondence in infants. One method is to project individuals’ electrode locations to a template brain with ROI parcellations constructed from a representative infant or an average template based on infant MRIs. Kabdebon et al. (2014) transformed infants’ (3 to 17 weeks) individual head models to a 7-week-old infant template and found stable correspondence between 10-20 electrode locations and underlying microanatomical ROIs. Electrodes O1, O2, T5 and T6 were projected to more inferior cortical regions than the cortical locations reported in Okamoto et al. (Okamoto et al., 2004). This study showed that age-related changes in the head and brain structures may lead to age-related variations in the correspondence between electrode positions and brain structures. Tsuzuki et al. (2017) transformed macroanatomical landmarks identified in 14 3- to 22-month-olds’ MRIs to a 12-month-old template (Matsui et al., 2014) in reference to virtual 10-10 electrode locations (Jurcak et al., 2007). They found inter-subject variations in the relative positions among the microanatomical landmarks. However, the differences were smaller than the region defined by the 10-10 electrodes. Hence, there may be relatively stable electrode-location-to-ROI correspondence across ages. One important limitation of these studies is the use of a single individual or single-age template as a reference for age ranges across which significant brain development occurs (e.g. 3 to 17 weeks; or 3 to 22 months).

The use of individual MRIs or age-appropriate MRI templates and stereotaxic atlases for standalone DOT data can reduce errors in co-registration and provide more accurate representations of electrode-location-to-ROI correspondence. Lloyd-Fox et al. (2014) co-registered channel locations (midway of between adjacent optode/electrode locations) on both the infants’ (4.5- and 6-month-olds) own MRIs and the age-appropriate average templates from the Neurodevelopmental MRI Database (Richards, in prep; Richards, Sanchez, Phillips-Meek, & Xie, 2015; Sanchez, Richards, & Almli, 2012a, 2012b). Anatomical stereotaxic atlases were constructed for individuals’ own MRIs and age-appropriate average templates (Fillmore, Richards, Phillips-Meek, Cryer, & Stevens, 2015). The correspondence between channel locations and ROIs in individual MRIs were highly comparable with the correspondence identified in the age-appropriate average MRI templates. An alternative to co-registering with the subjects’ own MRIs or average templates is to use an MRI from another infant with similar head size and age. A series of functional near-infrared spectroscopy (fNIRS) studies adopted Lloyd-Fox et al. (2014)’s co-registration method and positioned optode locations on close-head-size individual MRIs (Lloyd-Fox, Wu, Richards, Elwell, & Johnson, 2013) or both close MRIs and age-appropriate average templates (Emberson, Cannon, Palmeri, Richards, & Aslin, 2016; Emberson et al., 2017; Emberson, Richards, & Aslin, 2015). Spatial scalp projection was used to establish probabilistic mappings between channel locations and atlas ROIs. The channels with a large probability of localizing to a target ROI were used for group-level analyses.

### The Use of DOT Sensitivity to Describe Scalp-Cortex Correspondence

The use of spatial scalp projection for determining electrode-location-to-ROI correspondence is limited. It does not consider the interaction between near-infrared light and the optical properties of the biological tissues. Therefore, the approach is limited to spatially contiguous anatomical areas and cannot model directly the extent of the cortical regions being measured by NIRS recording nor the intensity of the signal generated by blood flow. An alternative to the spatial scalp projection is to use DOT sensitivity patterns to determine the correspondence between scalp recording locations and underlying cortical anatomy. DOT sensitivity represents the extent to which the DOT instrument can detect changes in brain activities in the region that it is sampling (Mansouri, L’Huillier, Kashou, & Humeau, 2010). It can be quantified as the fluence distribution at source location (“Direct DOT”) and the project of the fluence distribution at the source and the detector location (“Source-Detector Channel DOT”; Fu & Richards, under review).

The DOT sensitivity provides a measure of the scalp-location-to-ROI correspondence. The sensitivity distribution can be used to estimate the distance from scalp locations to the surface of cortical regions that are *measurable* with DOT and localize the ROIs that would be sampled with NIRS (Haeussinger et al., 2011). Studies have estimated DOT sensitivity in head models with atlas parcellations in infants (Bulgarelli et al., 2019; Bulgarelli et al., 2020; Perdue et al., 2019), children (Whiteman et al., 2017) and adults (Zimeo Morais, Balardin, & Sato, 2018). These studies computed the channel-ROI correspondence and generated look-up tables to show channel-ROI probabilities. Zimeo Morais et al. (2018) computed the specificity of each channel to the corresponding ROIs using the S-D Channel DOT measure. Such look-up tables facilitate the optimization of channel array design to maximize DOT sensitivity to user-specified ROIs (Brigadoi, Salvagnin, Fischetti, & Cooper, 2018; Zimeo Morais et al., 2018) and help to localize the possible ROIs that generated the fNIRS signals (Bulgarelli et al., 2019; Bulgarelli et al., 2020; Perdue et al., 2019).

### The Present Study

The current study examined age-related variations in the scalp-cortex correspondence. Our primary contribution was to use existing methods for DOT sensitivity across the period of infancy (2 weeks through 24 months) with narrow age bands, and compare these to older children (4, 12 years) and adults (20-24 years). Spatial scalp projection was used to estimate the distances between scalp electrode and channel locations and the cortical surface. We projected the electrode location to atlas locations delineated on individual MRIs to identify the anatomical mapping between scalp and ROI locations. We computed the distance between scalp locations and the cortical locations with maximum fluence. We provided a look-up table to present the probabilistic mapping between scalp channel locations and the underlying anatomical ROIs. Accurate and age-specific descriptions of scalp-to-cortex distance and scalp-to-ROI mapping are the bases for designing developmentally sensitive channel placements for NIRS recordings. Our look-up tables also provide important references for researchers to make anatomical interpretations of NIRS results from their specific age groups.

## Method

### Participants

The participants were 1058 typically developing participants ranging from 2 weeks to 24 years of age. The same sample was studied in Fu and Richards (under review). The MRIs were obtained from open-access databases and a local scanning facility. The sample consisted of 9 participants (4 females) from the Autism Brain Imaging Data Exchange (ABIDE; Di Martino et al., 2014), 280 (143 females) from the Baby Connectome Project (BCP; Howell et al., 2019), 177 (93 females) from the Early Brain Development Study (EBDS; Gilmore et al., 2010), 282 (106 females) from the Infant Brain Imaging Study (IBIS; e.g. Hazlett et al., 2017), 14 (5 females) from the Pediatric Imaging, Neurocognition, and Genetics Data Repository (PING; Jernigan et al., 2016), and 296 scans (141 females) from data collected at the McCausland Center of Brain Imaging (MCBI) or drawn from collaborative studies at other sites. Table 1 presents the number of MRIs for the open access databases, separately for age and gender. The sample ages were narrowest in the infancy period (2 weeks, 1-, 1.5-, 3-, or 6-month intervals from 2 weeks through 2 years) and included exemplar ages in children and adolescent ages (4, 12 years) and adult (20-24 years). All studies had institutional review board approval and informed consent for participants.

**Table 1.**
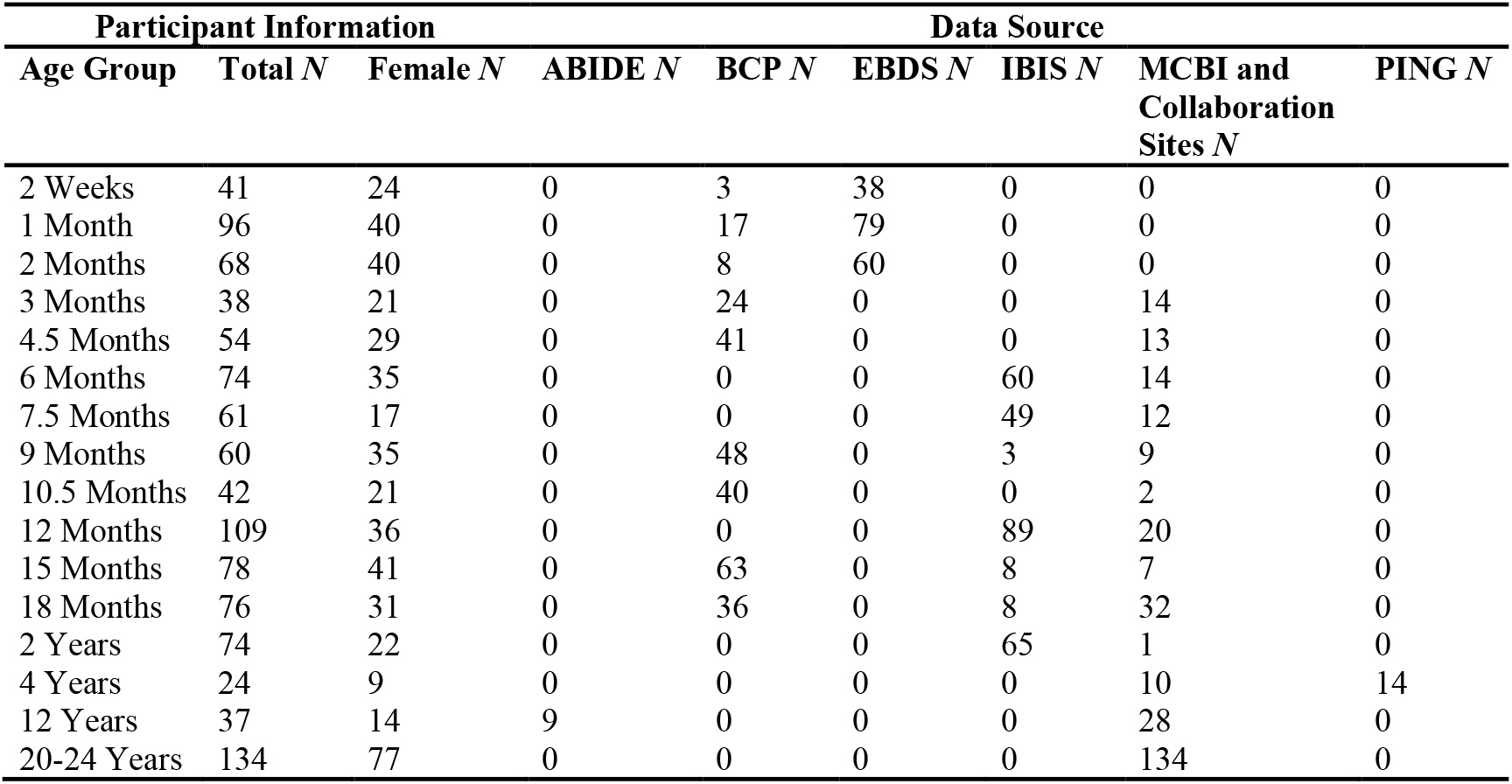
Demographical information of study participants by age group, sex, and data source. *Note*. ABIDE=Autism Brain Imaging Data Exchange; BCP=Baby Connectome Project; EBDS=Early Brain Development Study; IBIS=Infant Brain Imaging Study; MCBI=McCausland Center for Brain Imaging; PING=Pediatric Imaging, Neurocognition, and Genetics Data Repository

### MRI Sequences

The present study utilized T1-weighted (T1W) and T2-weighted (T2W) scans from each collection site. Details of the MRI acquisition protocols have been described in literatures on the Neurodevelopmental MRI Database (Fillmore, Phillips-Meek, & Richards, 2015; Fillmore, Richards, et al., 2015; Richards, in prep; Richards, Sanchez, et al., 2015; Sanchez et al., 2012a, 2012b). All MRIs were converted to NIFTI compressed format with 32-bit floating point resolution. Bias-field inhomogeneity correction (N4 algorithm) was performed on the extracted T1-weighted images (Avants et al., 2011; Tustison et al., 2010).

### MRI Preprocessing and Segmentation

First, the brains were extracted from the whole-head MRI volume in a procedure adapted from the FSL VBM pipeline (Douaud et al., 2007). The T1W volume for each participant was registered to an age-appropriate average MRI template. The MRI templates came from the Neurodevelopmental MRI database (Richards, Sanchez, et al., 2015; Sanchez et al., 2012a, 2012b). The brain from the average template was transformed into the participant MRI space and used a mask on the head volume. The extracted masked data was then used with the FSL brain extraction tool program (Jenkinson, Pechaud, & Smith, 2005; Smith et al., 2004). Each brain was visually inspected and manually modified if necessary. Second, each head MRI volume was segmented into 9 or 10 media types: gray matter (GM), white matter (WM), cerebrospinal fluid (CSF), non-myelinated axons (NMA), other brain matter, skin, skull, air, eyes, and other inside skull material. Details of the segmentation methods are presented in the Supplemental Information. The segmented regions were assembled into a single MRI volume we will refer to as the “segmented head MRI volume”. Figure 1A shows a 3D rending of the T1W volume from a 3-month-old infant with a cutout revealing the segmented MRI volume. The realistic head model represents the geometry of the head and allow us to differentiate optical properties of different tissue types.

**Fig 1.**
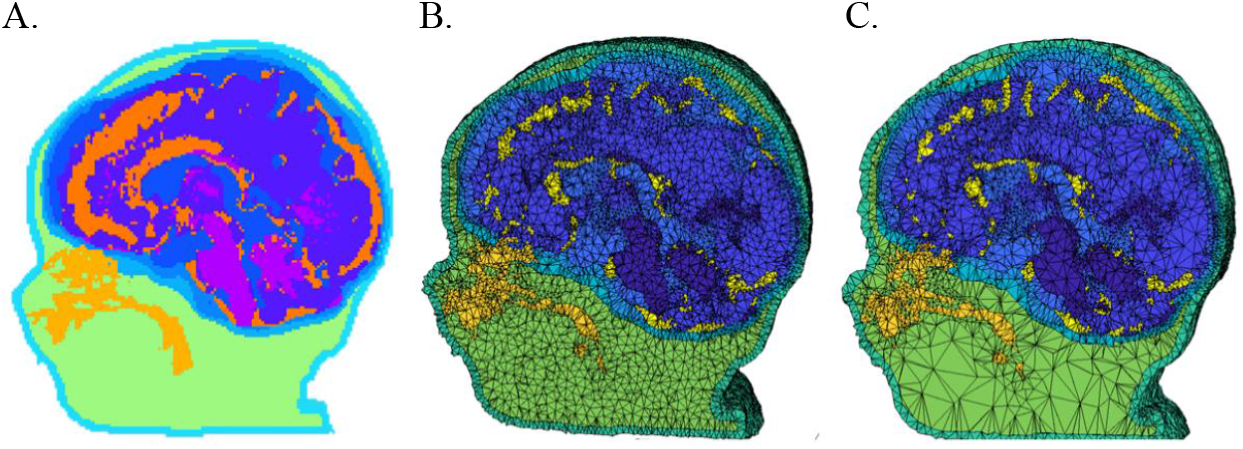
Segmented head MRI volumes. The examples were taken from the same 3-month-old infant MRI. A. The segmented head model. Aqua is the skull, purple is the gray matter, magenta is the white matter, dark orange is the cerebrospinal fluid, light blue is the dura, light orange is the nasal and mouth air cavities, and green is the non-bone structure (muscle, sinew, fat). B. The segmented head model with dense finite element model (FEM) mesh. C. The segmented head model with sparse FEM mesh.

### Mesh Generation

A “finite element method” (FEM) tetrahedral mesh was constructed from the segmented head MRI volume. Figure 1B and 1C displayed meshes that were produced using the iso2mesh toolbox with CGAL 3.6 mesh program (“v2m” function, Fang & Boas, 2009b). Tetrahedral meshes accurately represent boundaries of complex three-dimensional volumetric tissues and increase the accuracy in modeling photon propagation in the complex mediums such as the head and brain (Yan, Tran, & Fang, 2019). The FEM volumetric meshes have nodes that represent the voxel locations for the tetrahedra, a 4-element matrix representing the corners of each tetrahedron, and a vector representing the media type from the segmented head MRI volume. A mesh was generated for each segmented head MRI volume. Figure 1B depicts an example of the dense meshes. The number of nodes, elements, and tetra volume were calculated for the infants (2 weeks through 2 years), children (4 and 12 years), and adults (Supplemental Figure 1). We will refer to this as the “segmented FEM mesh”. We used the mesh to locate points on the scalp that were closest to the electrode positions, and for the segmented FEM mesh for the MMC computer program.

### Scalp Locations

#### Virtual Electrodes Placement

The locations for the 10-10 and 10-5 electrode systems were constructed on each head MRI volume. Figure 2 illustrates the placements of 10-10 and 10-5 electrodes positions. First, we manually marked cranial fiducial points using MRIcron (Rorden, 2012; Rorden & Brett, 2000): nasion (Nz), vertex (Vz), inion (Iz), left preauricular point (LPA), right preauricular point (RPA), left mastoid (LMa), and right mastoid (RMa) (Richards, Boswell, Stevens, & Vendemia, 2015). The coordinates of the fiducials were transferred on to the scalp mesh. Next, we calculated 81 virtual electrode positions based on the “unambiguously illustrated 10-10 system (Jurcak et al., 2007). Details for constructing the 10-10 locations are described in Richards, Boswell, et al. (2015) and the Supplemental Information. We simulated a total of 358 electrodes on 10-5 locations by calculating center points between 10-10 positions. The electrode positions also were computed for the average MRI templates.

**Fig 2.**
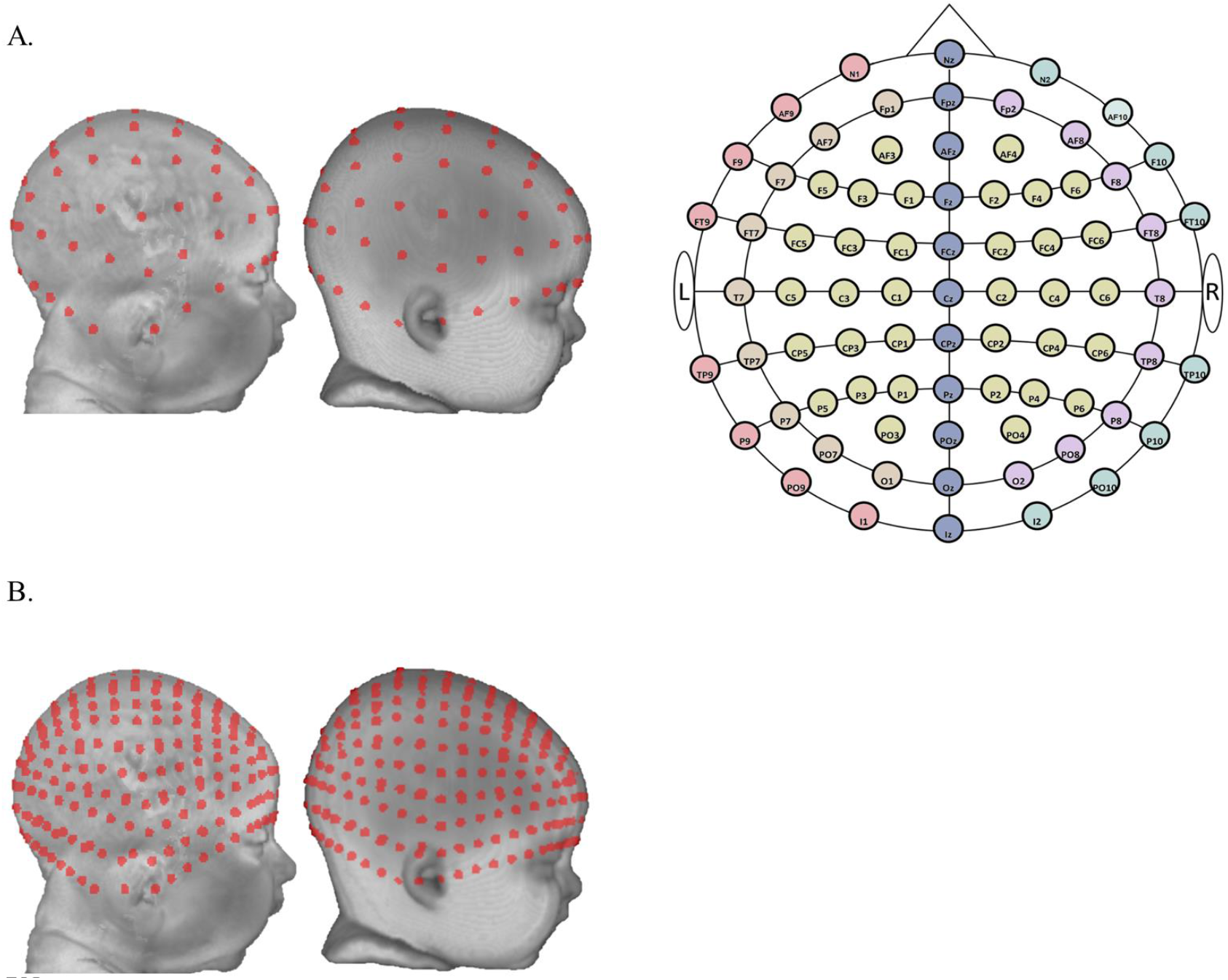
Virtual electrode placement. A. Ten-ten virtual electrode placement. From left to right: a three-month individual head model, an average template for three-month-olds, and a two-dimensional layout of the 10-10 system). Ten-ten electrodes were divided into six groups for visualization purposes. The six groups were color-coded on the two-dimensional schematic of the 10-10 system. Group 1: electrodes on the central curve (Nz-Cz-Iz). Group 2: electrodes on the left curve between Nz and Iz (N1-LPA/T9-I1). Group 3: electrodes on the right curve between Nz and Iz (N2-RPA/T10-I2). Group 4: electrodes on the left curve between Fpz and Oz (Fp1-T7-O1). Group 5: electrodes on the right curve between Fpz and Oz (Fp2-T8-O2). Group 6: the remaining electrodes enclosed by the central curve, the left and right curves between Fpz and Oz. B. Ten-five virtual electrode placement. From left to right: the same three-month individual head model and the three-month average template.

#### Source-Detector Channels

Source-detector channel locations were defined using electrode combinations centered on each 10-10 electrode. The 10-10 electrode locations were centered between surrounding adjacent pairs of 10-10 or 10-5 electrode locations. There were 251 source-detector pairs formed with adjacent 10-10 electrodes, *Mean*_Separation_ = 57.95 mm; *SD* = 14.22, and 251 pairs formed with adjacent 10-5 electrodes, *Mean*_Separation_ = 28.92 mm; *SD* = 7.06. The channel locations were used to estimate “S-D Channel DOT fluence” described below.

### Cortical Locations

Three stereotaxic atlases were constructed for each individual MRI. The atlases delineate cortical lobes or more specified locations within the lobes. We created a 1.5 cm spherical mask around each electrode or channel location to standardize the region to be identified in the atlases. The atlases and spherical masks were used with the spatial scalp projection method (described below) to describe the correspondence between scalp electrode positions and corresponding cortical structures in the individual’s own brain space with various spatial resolution (Lloyd-Fox et al., 2014). They were also used with the “S-D Channel DOT” estimates (described below) to identify the anatomical structures being sampled by the fluence distribution.

#### Stereotaxic Atlases

Three atlases were constructed for each individual MRI to delineate anatomical regions that can be used to identify brain locations corresponding to scalp electrode positions or DOT sensitivity patterns. Details of the atlas constructions may be found elsewhere (Fillmore, Richards, et al., 2015; Lloyd-Fox et al., 2014; Richards, 2013). The first atlas was the LONI Probabilistic Brain Atlas (LPBA40; Shattuck et al., 2008), which contains 56 areas from the cortical and subcortical regions, brainstem and cerebellum. The second was the Hammers atlas, based on MRIs from the IXI MRI database (Heckemann, Hajnal, Aljabar, Rueckert, & Hammers, 2006), consists of 83 areas defined from the cortex, subcortical, brainstem and cerebellum (Gousias et al., 2008). We used a majority vote fusion procedure that combines labeled segments from manually segmented MRIs to produce atlases that identify an anatomical area for each brain voxel of the individual MRI. The third atlas was an automatically constructed lobar atlas that identifies the major cortical lobes (e.g. frontal), some sublobar cortical (e.g. fusiform gyrus), subcortical (e.g. striatum), cerebellum and brainstem. The atlas was constructed by extracting and combining common areas from the LPBA40 and Hammers atlases, and manually segmented average template atlases transformed from the template space to the individual MRI space.

#### Spherical Masks

We created a sphere with 1.5 cm radius around each 10-10 electrode and channel location. Figure 3 shows examples of the spherical volumes were used as masks for determining the correspondence between scalp and brain locations. We identified the atlas regions of interest (ROIs) that intersected with the masks and computed the voxel numbers in the sphere of these ROIs. Hence, there was a distribution of atlas ROIs with various voxel numbers that were mapped to each electrode and channel location.

**Fig 3.**
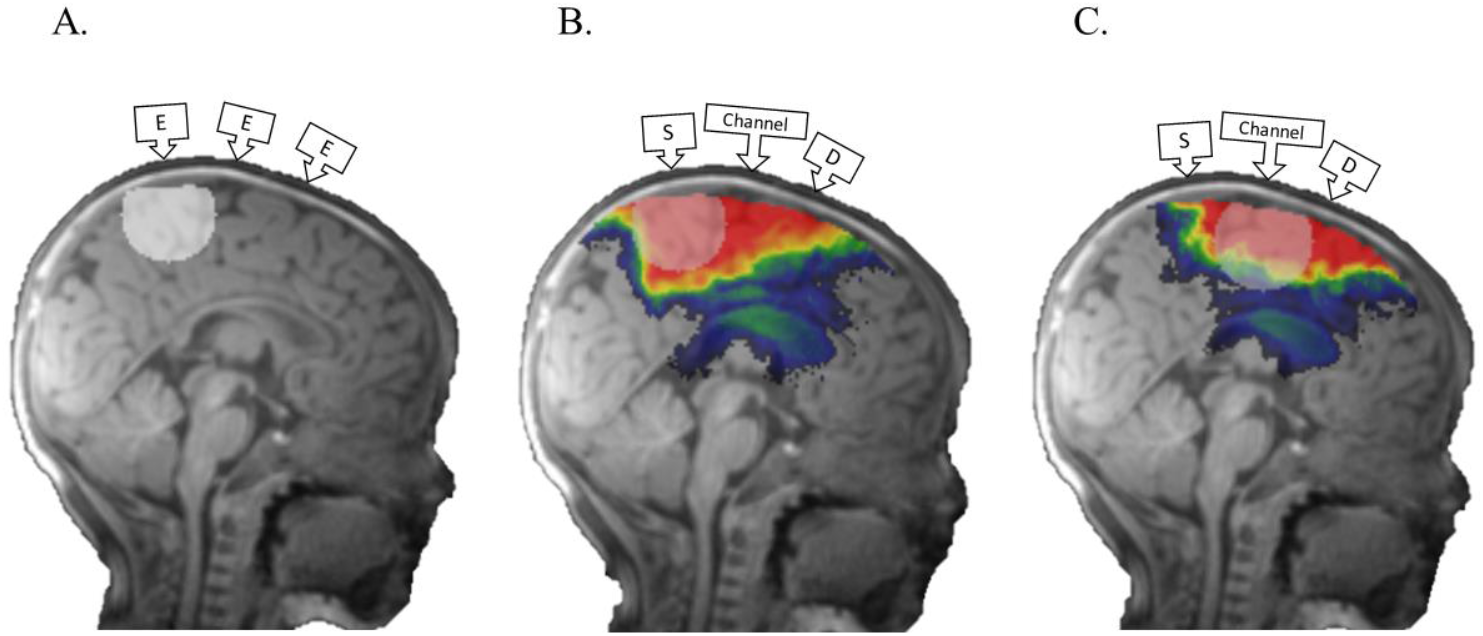
Methods for determining scalp-location-to-ROI mapping. The MRI was taken from a 3-month infant. A. An example of a 1.5cm-radius spherical mask created around an electrode location for the spatial scalp projection procedure. B. An illustration of a spherical mask created around a source location for the Direct DOT procedure. C. An illustration of a spherical mask created around a channel location for the Source-Detector (S-D) Channel DOT procedure. Monte Carlo photon migration simulations were used to estimate fluence distributions for individual 10-5 electrode locations (Direct DOT) and 10-10 channel locations (S-D Channel DOT). The red area represents greater fluence.

### DOT Sensitivity Analyses

The DOT fluence distribution was estimated from Monte Carlo simulations using the Monte Carlo eXtreme package (MCX; Fang & Boas, 2009a; Fang & Yan, 2019). Details of the simulations are presented in Fu and Richards (under review) and the Supplemental Information. The output from the Monte Carlo simulation contained the fluence across the entire MRI volume separately for each electrode. Figure 3B shows the DOT fluence plotted on an individual subject MRI for a single electrode location. We computed “S-D Channel DOT fluence” by multiplying the source-electrode (optode) fluence distribution by the detector-electrode (optode) fluence distribution. This represents the DOT fluence sensitivity for a photon channel from the source to the detector (“PMDF” in Brigadoi & Cooper, 2015). Figure 3C shows the S-D Channel DOT fluence plotted on an MRI for a single source-detector pair.

### Spatial Scalp Projection

We performed spatial projections from each 10-10 or 10-5 electrode position on the scalp to the brain surface. These projections were used to measure the distance between each scalp electrode and the brain surface and to examine electrode scalp location to anatomical ROI correlation (Emberson, Cannon, et al., 2016; Emberson et al., 2017; Emberson et al., 2015; Lloyd-Fox et al., 2014; Lloyd-Fox et al., 2013). The brain locations underlying each electrode was determined by an algorithm in which a spatial projection from the electrode scalp surface to the brain center was done. The point where the projection intersected the cortex was identified. These projections were done with the scalp and brain elements of the segmented FEM mesh and mesh manipulation tools from the iso2mesh toolbox (Fang & Boas, 2009b).

### Scalp-to-cortex Distance

We compared the distance from scalp electrode or channel locations to cortical surface across age groups. The “Scalp Projection” was simply the closest intersection of the brain from the projection of the electrode position toward the brain. The “Direct DOT” distance was based on the single-electrode DOT fluence. A 1.5 cm radius spherical mask was created at the point where the spatial projection intersected the brain. The maximum DOT fluence in the brain masked by the sphere was used as the location to calculate the Direct DOT distance. The “S-D Channel DOT” distance was based on the source-detector S-D Channel DOT fluence. The same spatial projection from the electrode, and the 1.5 cm sphere, was applied to the S-D Channel DOT fluence and the location of the maximum S-D Channel DOT fluence was used to calculate the S-D Channel DOT distance.

### Scalp-location-to-ROI Mapping

The Scalp Projection, Direct DOT and the S-D Channel DOT sensitivity were used to generate a look-up procedure that links the scalp electrode or channel locations to the lobar, Hammer, and LPBA40 atlas ROIs. A 1.5 cm mask was placed on the scalp projection to the brain intersection, maximum point of the DOT fluence, or maximum point of the S-D Channel DOT fluence. Figure 3A shows a sphere surrounding the scalp projection to the brain intersection; Figure 3B shows a sphere surrounding the electrode to the max DOT fluence location; Figure 3C shows a sphere surrounding the electrode to the S-D Channel DOT fluence. The anatomical ROI of each voxel in the respective spheres were recorded. The percentage of voxels in each ROI that intersected with the spherical masks was computed. We created tables for each age that listed the ROIs and the percentage of voxels in each ROI that intersected with the spherical masks created around the 10-10 electrode (Scalp Projection) and channel locations (S-D Channel DOT). The Scalp Projection look-up table details the spatial correspondence between the scalp electrode location and anatomical brain regions. The S-D Channel DOT look-up table illustrates the channel DOT sensitivity to cortical regions.

### Additional Measures and Analyses

There were several methods and results that are presented in the Supplemental Information. These include tMCimg (Boas et al., 2002) simulations in all individual MRIs and age-appropriate average templates and MMC simulations (Fang & Yan, 2019; Tran, Yan, & Fang, 2020; Yan et al., 2019) in 3-month and 6-month old individual MRIs and average templates. We compared the scalp-to-cortex distance estimations obtained from the MCX, tMCimg, and MMC simulation packages. *<u>fOLD Channels</u>*: We used a set of electrode pairs to define source-detector channels from the 130 channel locations specified in the fNIRS Optodes’ Location Decider (fOLD) (Zimeo Morais et al., 2018). We computed S-D Channel DOT fluence in selected age groups (3 month, 6 month, and 20-24 years) for the 130 channel locations using fluence estimated from MCX simulations (Zimeo Morais et al., 2018). We provided a look-up table that presented the specificity of each fOLD channel to the underlying ROI(s). We also compared scalp-to-cortex distances among simulation methods and brain model types.

## Results

### Scalp-to-Cortex Distance

The scalp-to-cortex distances were analyzed as a function of age for the Scalp Projection and S-D Channel DOT estimation methods. A one-way ANOVA for the infant ages (2 weeks through 2 years) was computed separately for the distances from the three estimation methods. There were significant main effects of age on the Scalp Projection, *F* (12, 856) = 19.01, *p* < .001, Direct DOT, *F* (12, 849) = 31.35, *p* < .001, and S-D Channel DOT distance measures, *F* (12, 849) = 31.86, *p* < .001. Figure 4 shows the average scalp-to-cortex distances for the age groups for the three estimation methods. The differences in the distances for the infant ages did not show a systematic age-related change. The Scalp Projection and Direct DOT distances were similar, and the S-D Channel DOT distances were larger than the distances from the other methods. We also tested whether scalp-to-cortex distances changed significantly from infancy (2 weeks to 2 years) to childhood (4 years and 12 years) and to adulthood (20-24 years). Results from the one-way ANOVAs confirmed the age group effect in distances estimated using scalp projection, *F* (2, 967) = 734.84, *p* < .001, Direct DOT, *F* (2, 1054) = 1708.43, *p* < .001, and S-D Channel DOT, *F* (2, 1054) = 1317.58, *p* < .001 (c.f. Figure 7B). The distances increased from infancy and toddlerhood to childhood and from childhood to adulthood for all estimation methods, all *p*’s < .001. Figure 4 shows an obvious change from the infant to the child/adolescent and adult ages in scalp-to-cortex distance.

**Fig 4.**
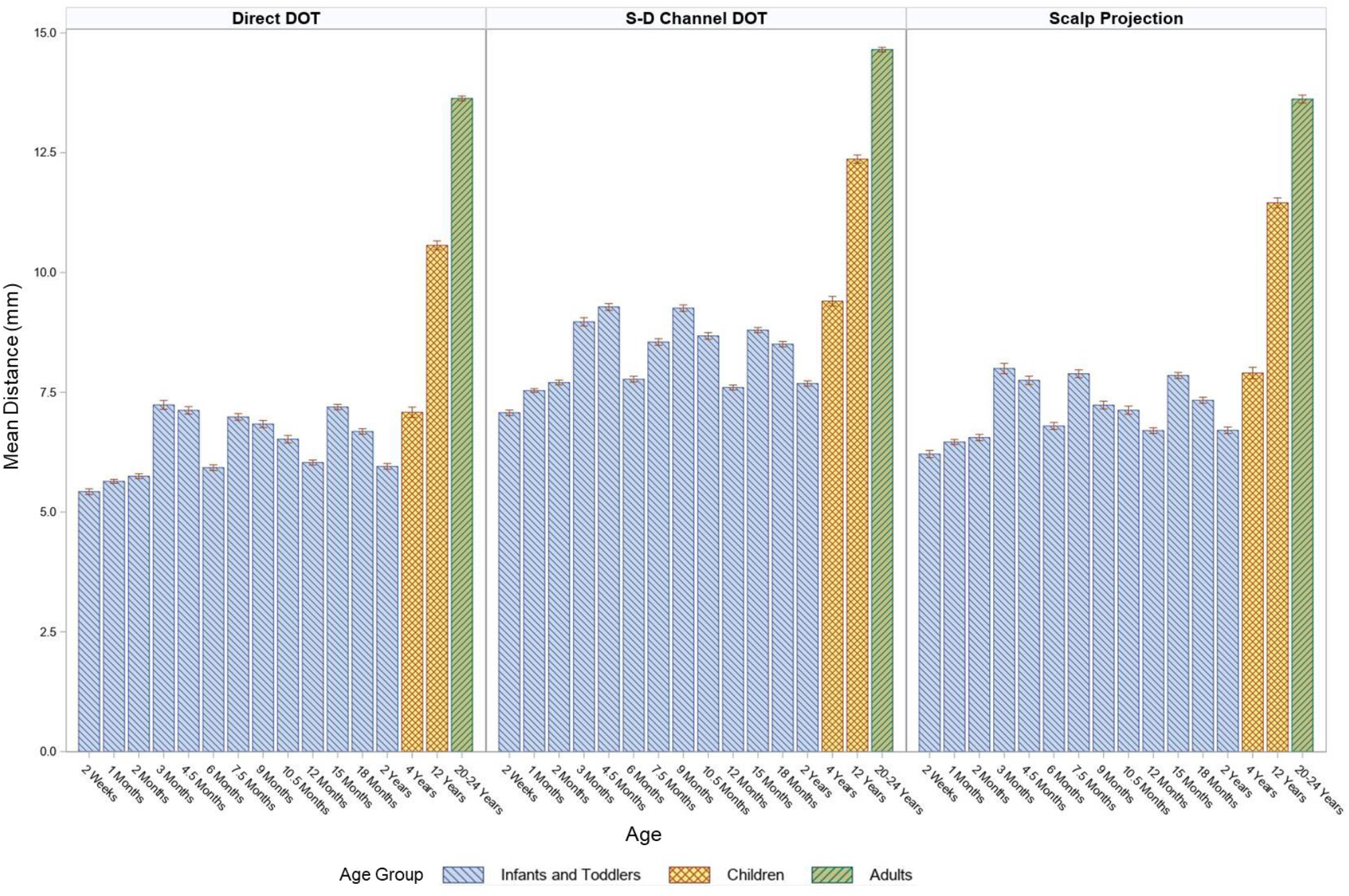
Mean scalp-to-cortex distances by age groups. The distances were estimated using scalp projection, Direct DOT and S-D Channel DOT with individual MRIs

We examined age-related changes in scalp-to-cortex distance. Figure 5 shows 2-D scalp topographical maps plotted with the Scalp Projection distance represented by the color map separately by age, and 3-D renderings of the distance on the head for three selected ages. There was an increase in the distance across electrode positions from the infants to the children to the adults. The electrodes on the bottom row (N1, AF9, F9, FT9, T9/LPA, TP9, P9, PO9, l1, N2, AF10, F10, FT10, T10/RPA, TP10, P10, PO10, and l2) showed the largest scalp-to-cortex distance, followed by the frontal and midline electrode positions, then followed by electrodes over the temporal, parietal and occipital positions. The shift from the frontal-midline electrodes to the other scalp positions was evident in several groups. Two-way ANOVAs with scalp positions as a within-subjects factor and age group as a between-subjects factor was conducted to examine age differences in distances by scalp locations among infant, child and adult age groups. We divided the electrodes to three groups based on Figure 5. Group one contained electrodes in the bottom row and midline. Group two contained frontal electrodes (‘F’ electrodes), and group three contained the rest of the electrodes. Among infants, there was a significant effect of age, *F* (12, 2547) = 64.28, *p* < .001, electrode group *F* (2, 2547) = 6936.95, *p* < .001, and age-by-electrode-group interaction effect, *F* (24, 2547) = 12.64, *p* < .001. The scalp-to-cortex distances of the group one electrodes were greater than those of group two and three electrodes (*p*’s < .001). The group two electrodes at frontal locations were more distant to the cortex than group three electrodes at central, temporal, parietal, and occipital locations for most of the infant groups, *p*’s < .01, except for 15 months (*p* = .25), 18 months (*p* = .23), and 2 years (*p* =.15). Among children, the main effect of age group, *F* (1, 177) = 260.89, *p* < .001, and electrode group, *F* (2, 177) = 406.58, *p* < .001, were significant. There was no age-by-electrode-group interaction effect, *p* = .022. The distance across all electrodes positions was greater in 12-year-olds than 4-year-olds, *p* < .001. The distance was largest in group one electrodes, *p*’s < .001, followed by group three electrodes, *p* = .025, across all child age groups. Among adults, the effect of electrode group was significant, *F* (2, 399) = 260.63, *p* < .001. Group one electrodes had the furthest distances to the cortex, *p*’s < .001, followed by group three electrodes, *p* = .002.

**Fig 5.**
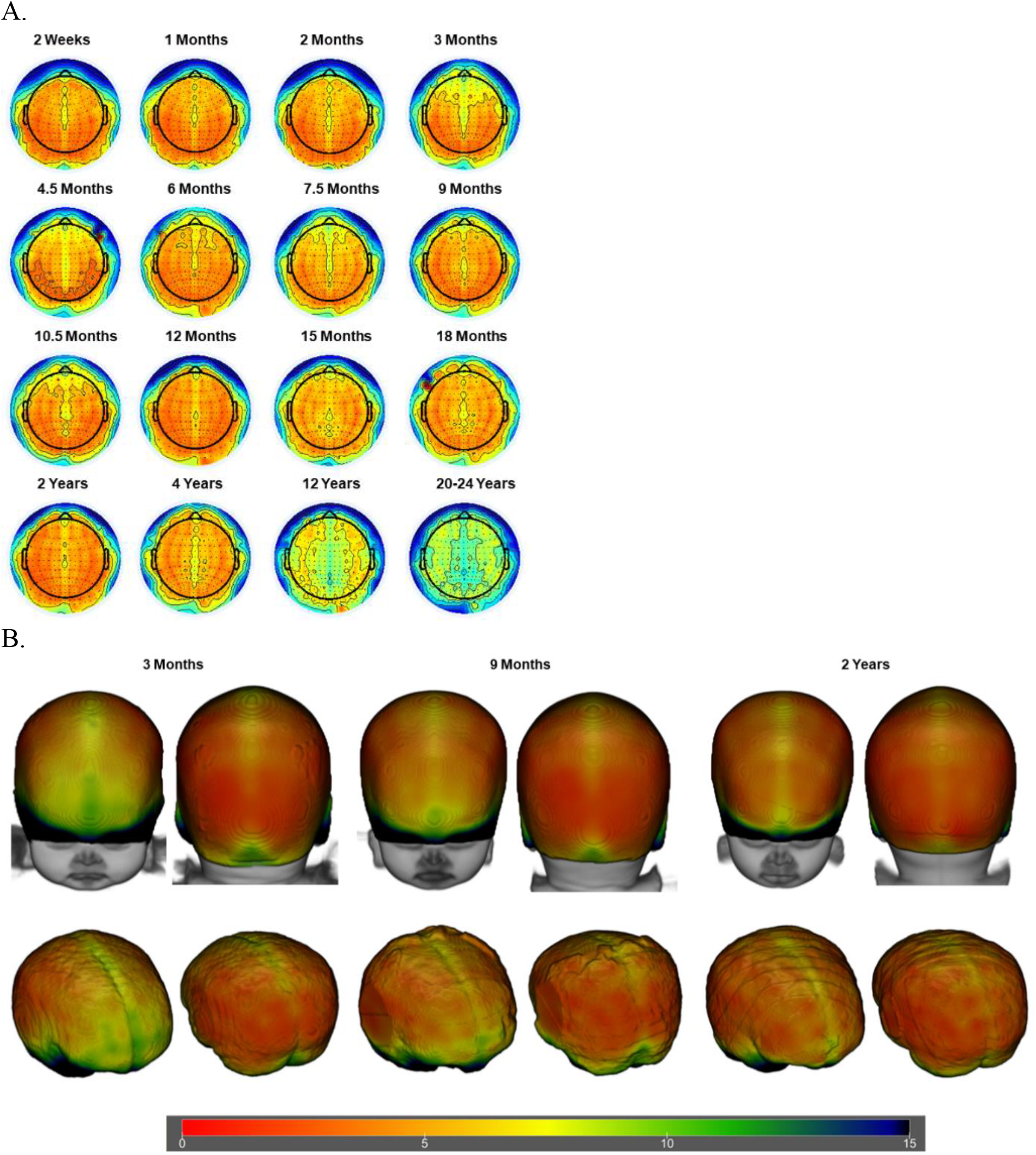
Mean distance from the scalp to cortical surface by electrode locations estimated using scalp projection. For the visualization purpose, estimations for the 10-5 electrode locations were displayed. The color bar denotes the distance range for all figure types. A. Scalp topographical maps for all age groups. Darker red represents closer scalp-to-cortex distances, and darker blue indicates greater distances. B. Three-dimensional rendering of mean distances on the heads (top row) and on the brains (bottom row) at 3-month, 9-months, and 2-years, selected as examples. Head and brain models were generated using age-matched average templates.

Figure 6 shows 2-D maps and 3-D renderings for the S-D Channel DOT scalp-to-cortex distance. The distance of the maximum S-D Channel DOT fluence was larger than the Scalp Projections distance at all ages (cf. Figure 5, 6). A similar pattern of distances across the scalp was found with the S-D Channel DOT distance. Among infants, there was a significant effect of age, *F* (12, 2547) = 66.27, *p* < .001, electrode group *F* (2, 2547) = 3635.67, *p* < .001, and age-by-electrode-group interaction effect on scalp-to-cortex distances, *F* (24, 2547) = 14.34, *p* < .001. The distances at group one electrode locations were the largest (*p*’s < .001). The distances at group two electrode locations were greater than group three locations for most of the infant groups, *p*’s < .05, except for 2 weeks (*p* = .92), 1 months (*p* = .49), 2 months (*p* = .74), and 18 months (*p* = .23). Among children, there was a significant effect of age *F* (1, 177) = 147.05, *p* < .001, and electrode group, *F* (2, 177) = 205.32, *p* < .001. The interaction effect was also significant, *F* (2, 177) = 3.67, *p* =.03. Group one electrodes had the furthest distances to the cortex for both 4- and 12-year-olds, *p*’s < .001. Distances at group two electrode locations were greater than group three locations for 12-year-olds, *p* = .02, but not for 4-year-olds, *p* = .29. Among adults, the effect of electrode group was significant, *F* (2, 399) = 280.34, *p* < .001. Group one electrodes had the furthest distances to the cortex, *p*’s < .001, followed by group three electrodes, *p* = .001. The findings indicate that the scalp-to-cortex distances were largest in bottom-row and midline electrodes for all age groups. The anterior-to-posterior decreases in scalp-to-cortex distance were visible in most of the infant groups.

**Fig 6.**
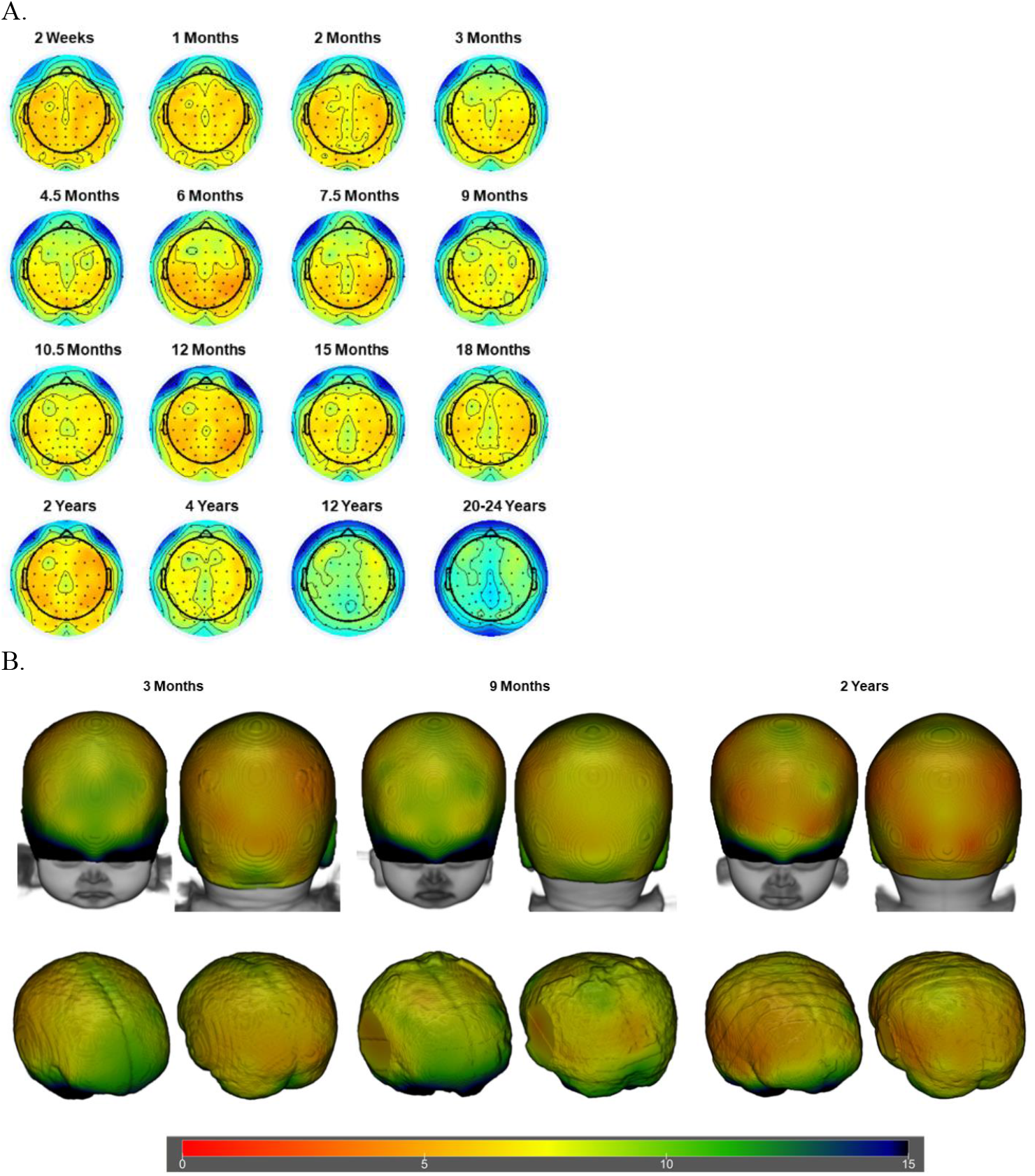
Mean S-D Channel DOT distances at 10-10 channel positions. The color bar denotes the distance range for all figure types. A. Scalp topographical maps for all age groups. Darker red represents closer scalp-to-cortex distances, and darker blue indicates greater distances. B. Three-dimensional rendering of mean distance on the heads (top row) and on the brains (bottom row) at 3-month, 9-months, and 12-years, selected as examples. Head and brain models were generated using age-matched average templates.

We inspected changes in the scalp-to-brain distances across the individual 10-10 electrode positions. The distance measures were averaged across three age groups (infants and toddlers: 2 weeks to 2 years; children: 4 years and 12 years; and adults: 20-24 years), electrode position, and estimation methods (Scalp Projection, Direct DOT, and S-D Channel DOT). Figure 7A shows the scalp-to-cortex distances for the 10-10 positions separately for the three age groups and the three estimation methods. Two-way ANOVAs were conducted to examine the effect of estimation method, electrode group (six groups based on Figure 7), and interact effect on mean distances among the infant, child, and adult age groups. The main effect of estimation method was significant for infants, *F* (2, 15500) = 1177.97, *p* <.001, children, *F* (2, 1068) = 60.72, *p* <.001, and adults, *F* (2, 1890) = 39.91, *p* <.001. The main effect of electrode group was also significant for infant, *F* (5, 15500) = 11697.60, *p* <.001, children, *F* (5, 1068) = 521.70, *p* <.001, and adults, *F* (2, 1890) = 591.76, *p* <.001. The interaction effect between estimation method and electrode group was significant for infants, *F* (10, 15500) = 102.36, *p* <.001, children, *F* (10, 1068) = 4.14, *p* <.001, and adults, *F* (10, 1890) = 2.08, *p* <.02. The larger distances were at bottom row electrode position for all estimation methods and age groups, *p*’s <.001. This was followed by the midline electrodes and then by the electrodes in the other positions for all estimation methods and age groups, *p*’s <.001. Figure 7B displays the mean S-D Channel DOT distances for the infant, child, and adult group averaged across the 81 channel positions and individual age groups. We also found that the distances for the S-D Channel DOT fluence for the participant-based averages (e.g., Figure 7A) were similar to the distances calculated from the average MRI template for the same age (Supplemental Information Figure 3A).

**Fig 7.**
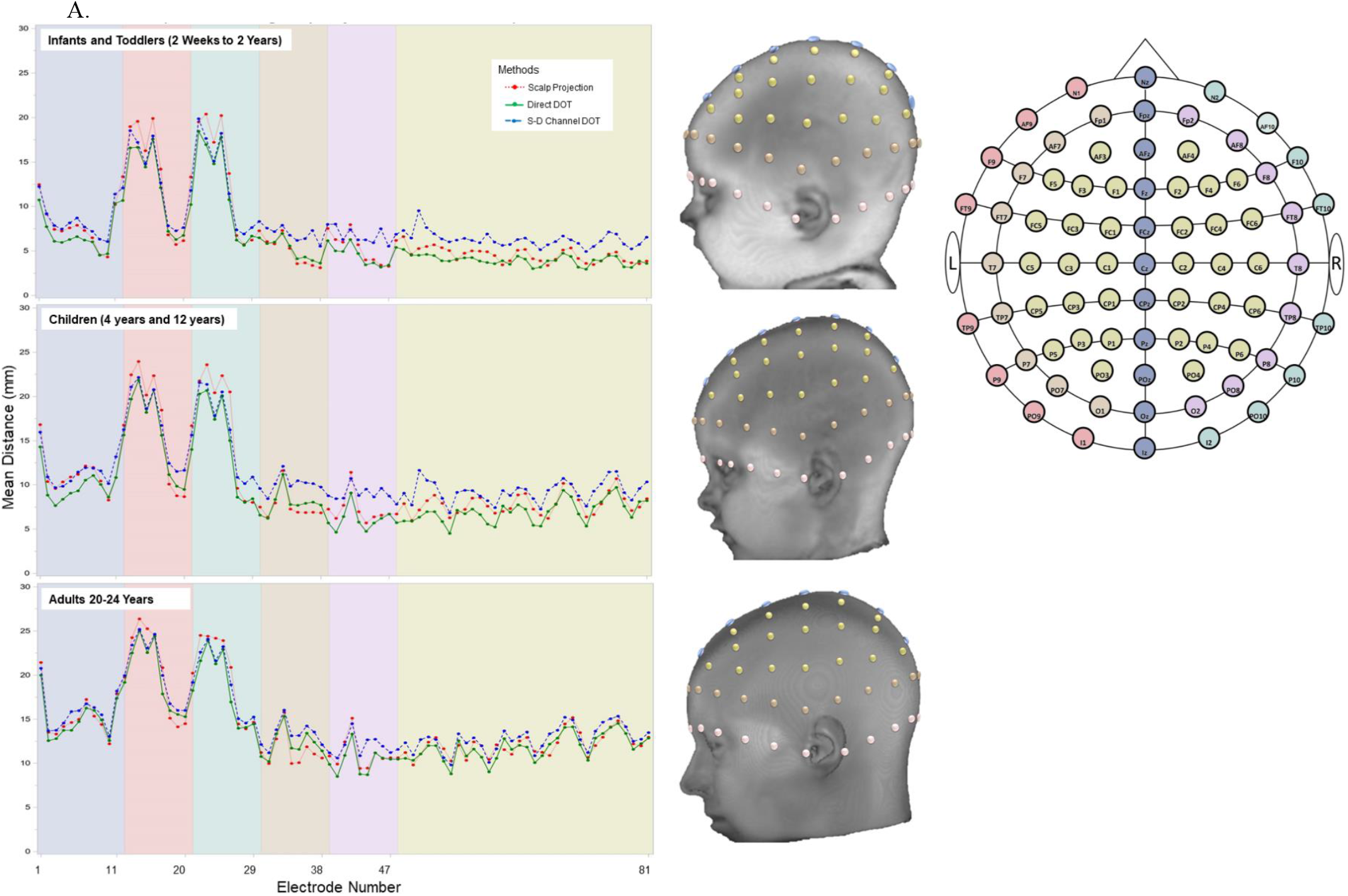
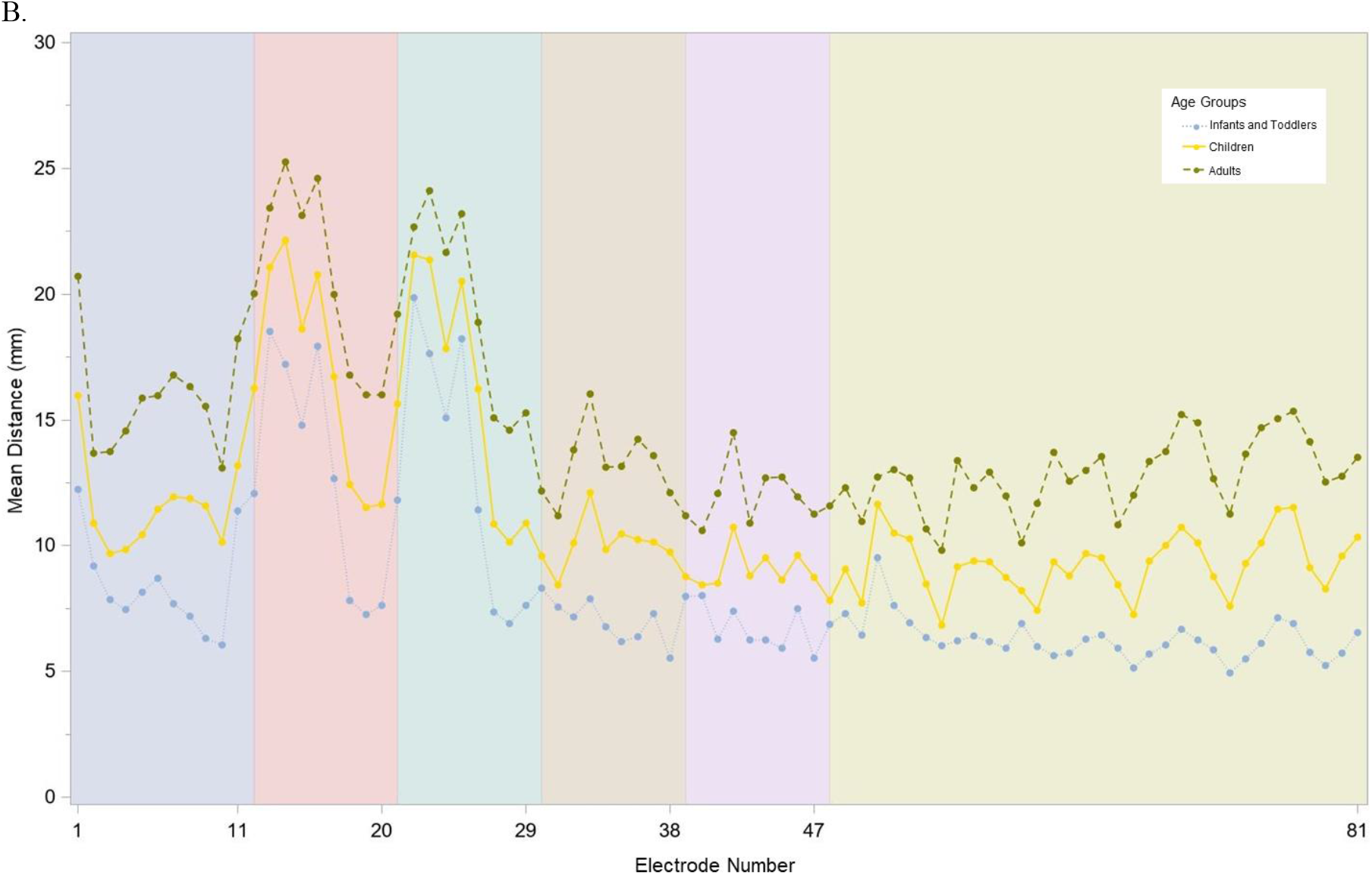
Mean scalp-to-cortex distances across 10-10 electrode and channel locations by age groups. A. Distances estimated using Scalp Projection, Direct DOT and S-D Channel DOT with individual MRIs. Electrodes were divided into five groups. In addition, a two-dimensional layout of the 10-10 system (top right), a three-dimensional rendering of the electrode locations on a 3-month-old infant average template (top left), a 4-year-old child average template (middle) and a 20-24-year-old adult average template (bottom) were displayed to aid visualization. The 10-10 electrode name and the number in the figure are in Table 3, and the five electrode groups are listed in Figure 1. B. Mean S-D Channel DOT distances for infants and toddlers (2 weeks to 2 years), Children (4 years and 12 years) and adults (20-24 years).

The scalp-to-cortex distances were examined also for the average MRI templates. Supplemental Figure 3 shows the S-D Channel DOT distance across age for the distances calculated from the average MRI templates. The distances for the participant-based averages (e.g., Figure 4, Figure 7B) were similar to the distances calculated from the average MRI template (Supplemental Information Figure 3B).

### Scalp-location-to-ROI Mapping

The Scalp Projection and the S-D Channel DOT sensitivity were used to generate look-up tables that show the cortical ROIs for each of the 10-10 electrodes. An Excel spreadsheet is presented in the Supplementary Information with this information (Supplemental Information: Table1). This table has a tab for each estimation type and age combination (e.g., Spatial 2-0 Weeks, Spatial 1-0Month, Source-Detector-DOT 2-0Weeks, Source-Detector-DOT 1-Month….). Each table has one row for each electrode containing columns for the electrode name and the cortical ROIs for the lobar, Hammers, and LPBA40 atlases. This table could be used to find a specific electrode-ROI combination for each of the ages in the study. We use the name “electrode” and “channel” interchangeably, though the “electrode location” properly refers to the Scalp Projection method where the “channel location” properly refers to the S-D Channel DOT method.

The Scalp Projection and S-D Channel DOT estimations yielded overlapping and considerably different scalp-location-to-ROI Mappings. For example, between-method discrepancy was found at 32 scalp locations where the electrode was mapped to at least one different ROI for the 3-month group. The discrepancy was found at 29 scalp locations for the 20-24-year-olds.

The channel-location-to-ROI correspondences showed stability over age for some of the electrode/atlas ROIs based on the S-D Channel DOT sensitivity estimation. The lobar atlas has macrostructural segmentations which resulted in similar channel-to-ROI correspondence across ages. Most of the channel locations (52 out of 81 channels) were mapped to one lobar atlas ROI for all age groups. Some channels were sensitive to more than one lobar ROIs in younger age groups but were mapped to one lobar ROI in older age groups (CZ, PZ, OZ, F9, F10, T10, FT7, FT8, O1, O2, C5, C6, C1, C2, CP5, and CP6). For example, O1 (left) and O2 (right) were mapped to the cerebellum, fusiform gyrus, occipital lobe, and parietal lobe for the infant groups but only to the occipital lobe for 20-24-year-olds. Some channels were sensitive to more than one lobar ROIs only in older age groups (POZ, PO9, I1, and I2). For example, I2 corresponded to both the right cerebellum and occipital lobe in 12- and 20-24-year-olds but only to the cerebellum in younger ages. Other channels had similar channel-to-ROI mappings over age (IZ, P7, P8, PO7, PO8, C3, C4, PO3, and PO4).

The Hammers and LPBA40 atlases have smaller structural segmentations than the lobar atlas. Hence, most of the channel locations were mapped to more than one ROI. At 44 channel locations, at least one identical ROI from the Hammers and LPBA40 atlas was mapped to the channel locations across all age groups. There were channels that were sensitive to superior frontal gyrus (FPZ, AFZ, FZ, FCZ, F2, FC1, and FC2), middle frontal gyrus (FP1, FP2, AF3, AF4, F5, F3, F4, F6, FC3, and FC4), inferior frontal gyrus (F7, F8, F5, F6, FC5, and FC6), precentral gyrus (FC5, C1, and C2), postcentral gyrus (C3 and C4), superior parietal gyrus (CPZ, CP1, and CP2), angular gyrus (CP3 and P4), cuneus (POZ), inferior temporal gyrus (FT9, T9, and T10), middle temporal gyrus (T7 and T8), and cerebellum (TP9, P9, PO9, I1, TP10, P10, PO10, and I2) across all age groups. Many channel locations displayed overlapping channel-to-ROI mapping while between-age differences were also observed. Figure 8 provides examples of the channel locations sensitive to ROIs delineated using LPBA40. Figure 8A shows a consistent pattern of channel locations across age for the inferior frontal gyrus, AF7, F5, F7, FC5, AF8, F6, F8, and FC6; channel location F3 also was sensitive to the inferior frontal gyrus at age 12 months. Figure 8B shows a less consistent pattern of channel locations sensitive to the postcentral gyrus, including CPz, CP1, C1, C3, C4, C5, and C6. However, the pattern of sensitive channel locations was different for each age.

**Fig 8.**
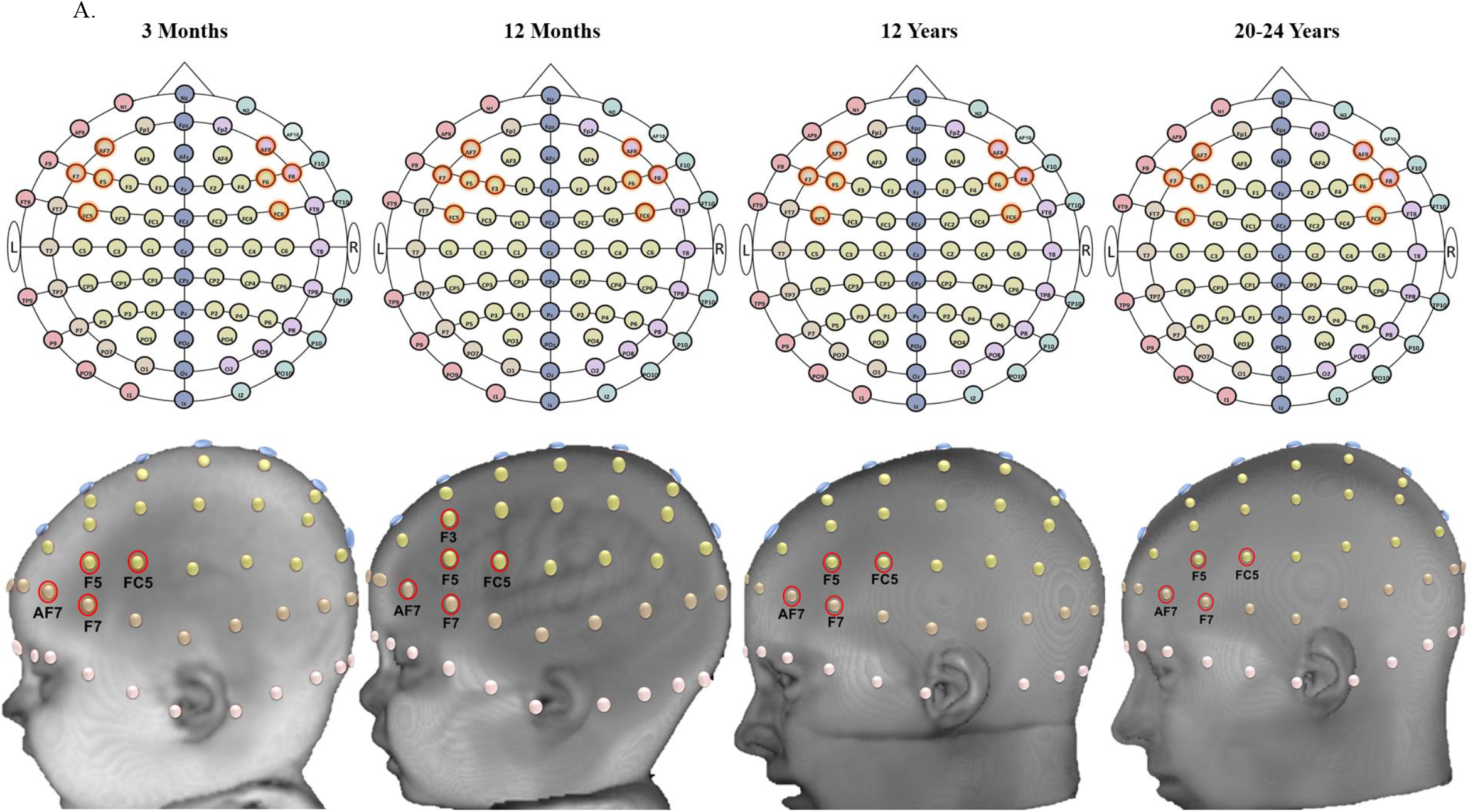
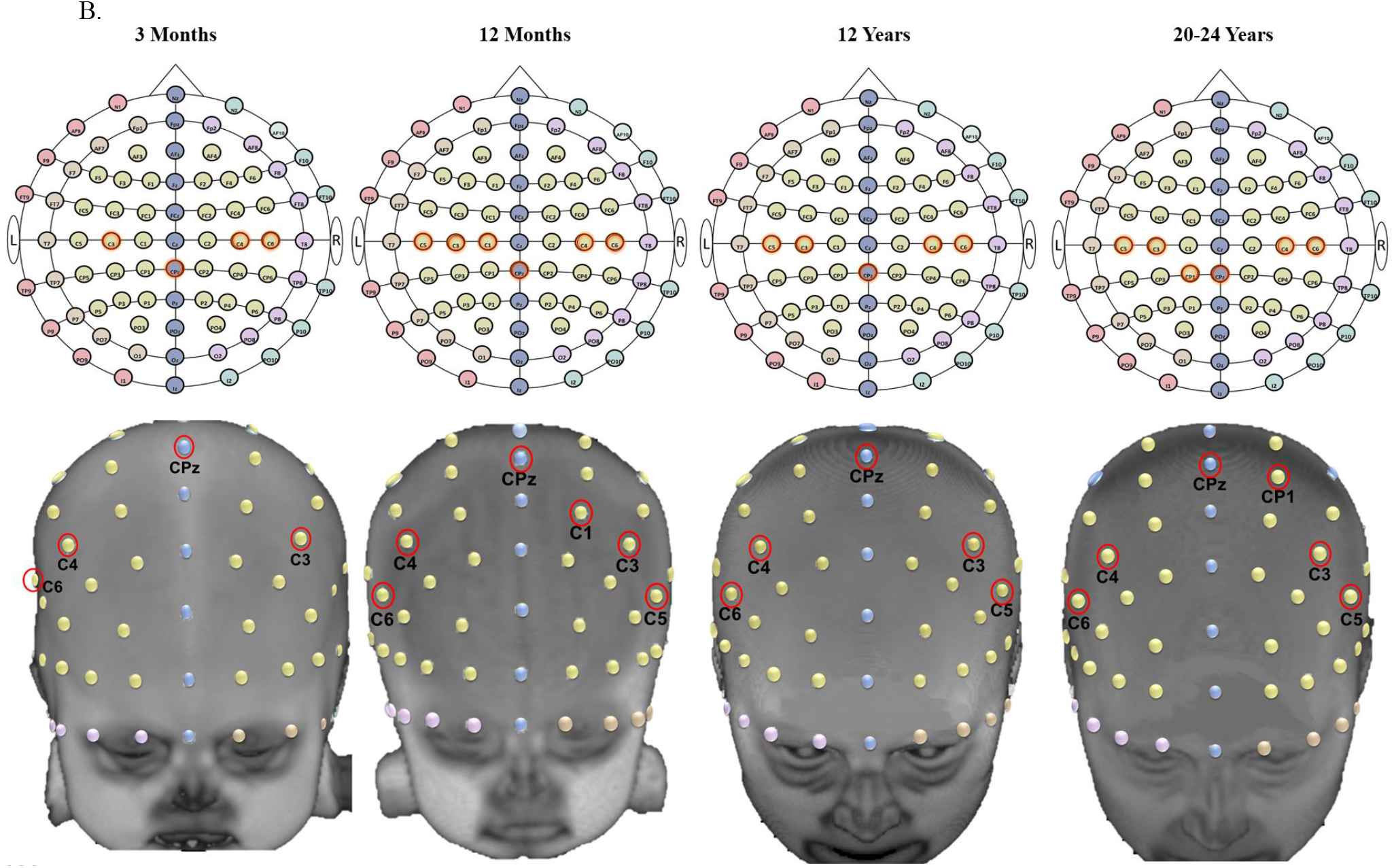
Examples of scalp-to-ROI mapping using S-D Channel DOT estimation in selected age groups (3 months, 12 months, 12 years, and 20-24 years). ROIs are delineated using the LONI Probabilistic Brain Atlas (LPBA40; Shattuck et al., 2008). Channel locations are displayed on age-matched average templates. A. S-D channel locations sensitive to the inferior frontal gyrus (the left inferior frontal gyrus is shown on the head model). B. S-D channel locations sensitive to the postcentral gyrus.

## Discussion

The present study examined age differences in scalp-to-cortex distances and scalp-location-to-ROI correspondence. We extended existing co-registration and photon migration simulation methods to realistic head models from infants ranging from 2 weeks to 24 months, children (4, 12 years) and adults. The scalp-to-cortex distances increased from infancy to childhood and to adulthood. There were considerable variations in the distance measures among the infant age groups. The probabilistic mappings between scalp locations and cortical lobes were relatively stable across development. However, the mappings between scalp locations and cortical ROIs in sub-lobar atlas parcellations showed greater age-related variations. We found that individual participant MRIs and average MRI templates from the Neurodevelopmental MRI Database (Richards, in prep; Richards, Sanchez, et al., 2015; Sanchez et al., 2012a, 2012b) were similar in their scalp-to-cortex distances.

### Scalp-to-Cortex Distance

Three methods were used to measure scalp-to-cortex distances. The Scalp Projection method has been commonly adopted in developmental and adult studies to measure the anatomical distance between scalp electrode locations and the cortical surface (Beauchamp et al., 2011; Emberson et al., 2017; Kabdebon et al., 2014; Okamoto et al., 2004; Whiteman et al., 2017). We also estimated the distance between the scalp location and the maximum source-detector fluence for the “Direct DOT” fluence distribution. The distances for these two measures were very similar. This is likely due to the monotonic and somewhat non-linear photon decay distribution across the head which is maximal near the surface of the cortex (Fu & Richards, under review). The S-D Channel DOT fluence distances were greater than distances estimated from the other methods across scalp locations for all age groups. The S-D Channel DOT fluence represents the flow of photons from the source to the detector (Fu & Richards, under review) and was deeper than the cortical surface (Scalp Projection) or the photon flow from the optode source injection point (Direct DOT).

All three estimation methods indicated that scalp-to-cortex distances averaged across the scalp locations increased from infancy to childhood and from childhood to adulthood. This finding is consistent with Beauchamp et al. (2011) who measured whole-brain distances from the scalp in newborns through 12-year-olds and other studies that measured scalp-to-cortex distances by electrode positions (3.4- to 16.3-week-olds; Kabdebon et al., 2014; 22- to 51-year-olds; Okamoto et al., 2004; 5- to 11-year-olds; Whiteman et al., 2017). The increase in scalp-to-cortex distances from infancy to adulthood may be attributed to several types of changes in the structure of the head. These include skull thickness (Emberson, Crosswhite, Goodwin, Berger, & Aslin, 2016; Hansman, 1966), increases in CSF volume (Makropoulos et al., 2016), global brain volume and gray matter / white matter growth (Gilmore et al., 2011; Makropoulos et al., 2016; Richards & Conte, 2020; Richards & Xie, 2015), and cortical folding during the first 2 years (Li et al., 2014). Brain structural growth is more gradual during childhood and adolescence and reaches plateau during adulthood (Brain Development Cooperative Group, 2011; Raznahan et al., 2011; Richards & Conte, 2020; Richards & Xie, 2015).

There were between-electrode variations in scalp-to-cortex distance, and these variations were heterogeneous among infant groups. The Scalp Projection and S-D Channel DOT distances for all age groups were greatest at the frontal and central electrodes on the bottom row around the face area, followed by electrode locations at the midline along the intra-hemispherical fissure, and the smallest on the electrode locations internal to these edges. Such inter-electrode variations were also found in previous studies (Kabdebon et al., 2014; Okamoto et al., 2004; Whiteman et al., 2017). Furthermore, we extended prior findings with young infants (Kabdebon et al., 2014) and revealed that there was a prominent decrease in distances from anterior to posterior locations in some infant groups (Scalp Projection distances in 3- to 10.5-month-olds and S-D Channel DOT distances in 2-week- to 10.5-month, 15-month, and 18-month-olds). This anterior-to posterior decrease in distance was less discernible in other age groups including adults (Okamoto et al., 2004). The anterior-posterior gradient coincides with the posterior-to-anterior sequence of cortical maturation (Brain Development Cooperative Group, 2011; Gilmore et al., 2007). The occipital and parietal lobes show faster growth in volume during infancy than the frontal regions (Gilmore et al., 2007). This implies that the posterior regions have expanded closer to the scalp surface earlier than anterior regions. In contrast, there is more significant age-related decline in occipital and parietal lobes than the frontal lobe from childhood to young adulthood (Brain Development Cooperative Group, 2011). The region-specific heterogeneous patterns of brain development that are observable from infancy (e.g. Gilmore et al., 2007; Gilmore et al., 2011; Li, Lin, Gilmore, & Shen, 2015; Makropoulos et al., 2016; Remer et al., 2017) may contribute to the lack of systematic age-related increase in scalp-to-cortex distances among infant groups. Together, the Scalp Projection and DOT fluence measures of scalp-to-cortex distances are sensitive to the complex age- and region-dependent cortical growth.

Gauging scalp-to-cortex distance is a foundational step for optimizing DOT sensitivity to the target cortical regions. Fu and Richards (under review) showed that the infants groups displayed different DOT sensitivity profiles compared to the adults with source-detector channels placed at the same distances. A common practice is to use longer separations for adults than infants (Gervain et al., 2011; Pinti et al., 2020). Increasing the separation distance allows the fluence distribution to extend deeper into the head tissues and thus sample cortical regions at greater depth (Fu & Richards, under review; Strangman, Li, & Zhang, 2013). However, the increased depth sensitivity is at the expense of decreased signal strength (Wang, Ayaz, & Izzetoglu, 2019). The current findings on age differences in scalp-to-cortex distance can be used with age-specific DOT sensitivity profiles (Fu & Richards, under review) to find optimal source-detector separation distances that allow for comparable depth sensitivity for different age groups.

Age-appropriate MRI templates may be used to describe cranio-cerebral correspondence when subjects’ own MRIs are unavailable. The present study showed that the age-appropriate templates has comparable S-D Channel DOT distances as the individual head models. Furthermore, the S-D Channel DOT distance estimations were robust across different Monte Carlo simulation methods (MCX, MMC, and tMCimg) for 3-month and 6-month infants’ individual MRIs and age-appropriate average templates (Supplemental Information Figure 5, 6, and 7). The developmental differences in scalp-to-cortex distances suggest that adult head models should not be used to make anatomical inferences for infant or child standalone NIRS/fNIRS data. We recommend using age-appropriate average templates or the individual MRIs when the subject’s own MRIs are not collected (e.g. Emberson, Cannon, et al., 2016; Emberson et al., 2017; Emberson et al., 2015).

### Scalp-location-to-ROI Mapping

This is the first study that provided scalp-location-to-ROI look-up tables computed using multiple methods for a wide range of age groups. We used the Scalp Projection and the S-D Channel DOT fluence to map 10-10 scalp electrode/channel locations with underlying ROIs from a macrostructural atlas (lobar) and two sublobar atlases (Hammers and LPBA40). The Spatial Projection method mapped underlying ROIs based only on the spatial anatomical relations. The S-D Channel DOT method mapped underlying ROIs based on the DOT fluence sensitivity profile. The S-D Channel-DOT look-up table supported the prior finding that there is consistent correspondence between the majority of scalp locations and macrostructural ROIs for infants (Kabdebon et al., 2014; Lloyd-Fox et al., 2014) and adults (Koessler et al., 2009; Okamoto et al., 2004). For smaller sub-lobar atlas ROIs, some channel locations show consistent correspondence between scalp location and underlying cortical ROI. For example, F5 and F6 were corresponded to bilateral inferior and middle frontal gyrus in all age groups. However, many channel locations show inconstant mappings across age groups. These include posterior midline positions and frontal channels on the bottom row where the scalp-to-cortex distance was larger. For example, channel TP7 was sensitive to both the inferior and middle temporal gyrus in the LPBA40 atlas for age 2 weeks through 12 years whereas it was mapped to only the middle temporal gyrus for 20-24-year-olds. Therefore, it is important to use age-appropriate head models to fully account for the age-dependent head and cortical structural changes when examining cranio-cerebral correspondence.

The age-appropriate S-D Channel DOT look-up procedures can be adopted to localize the ROI(s) that generate the fNIRS activities and design the optode arrangement prior to data collection. Our findings highlight a recognized issue in fNIRS data interpretation: the channel(s) that show significant activations from the group analysis may not correspond to the same ROI for all participants (Powell, Deen, & Saxe, 2018). This problem is especially concerning for infant and child studies that encompass a wide age range. Our S-D Channel DOT look-up procedures provide an effective solution. The optode locations recorded from an experiment can be co-registered with individual head models from infants closely matched in age and head measurements, if subjects’ own MRIs are not available. S-D Channel DOT sensitivity is then estimated to infer the subject-specific ROI(s) that have generated the fNIRs signals. The methodological details were also presented elsewhere (Bulgarelli et al., 2019; Bulgarelli et al., 2020; Perdue et al., 2019).

Our method and tables may also be used to design optode placement on NIRS holders that maximize channel sensitivity to hemodynamic changes of underlying ROIs. The tables we provide for the 10-10 or 10-5 recording system can be used as lookup tables for either manual construction of optode locations or used with automatic methods. We additionally provided age-appropriate S-D Channel DOT look-up table that quantify the sensitivity of channel pairings from the fOLD toolbox (Zimeo Morais et al., 2018) to lobar and sub-lobar ROIs (Supplemental Information Table 2). The table provides the specificity (%) of the channel to the ROI out of the total S-D Channel DOT sensitivity to all ROIs in the atlas. Researchers can thus design their holders to include channels that are sensitive to the user-specified ROIs with a specificity threshold (Zimeo Morais et al., 2018). Supplemental Information Figure 2D presents an example of optode placement that includes channels sensitive to the left inferior frontal gyrus with specificity greater than 1% for 6-month infants.

### Implications

The age-specific scalp-to-cortex distances and channel-to-ROI look-up tables can be used to guide channel placements and data analysis. Cross-sectional and longitudinal fNIRS studies need to ensure that age-group comparisons are made on data from sensors that sample the same ROI with comparable sensitivity to the cortex across ages. For example, for a study aiming to compare activations in the inferior frontal gyrus among 3-month, 12-month, and 12-year-old participants, Supplementary Information Table 1 and Figure 8A indicate that sensor placement that covers channel locations AF7, F7, F5, F3, FC5, AF8, F8, F6, and FC6 could be used for all age groups. We also know that the average distances to the cortex from the frontal electrode locations differed by age (3 months: 8mm; 12 months: 6mm; 12 years: 10mm). Hence, larger source-detector separation distances are expected for older ages. fNIRS studies conventionally use 20mm to 30mm separations for infants (Gervain et al., 2011; Lloyd-Fox, Blasi, & Elwell, 2010) and 30mm to 35mm for children (e.g. Ding, Fu, & Lee, 2014; Perlman, Luna, Hein, & Huppert, 2014). Experimenters should measure the source-detector separation distances after constructing a preliminary holder and adjust accordingly. Experimenters are not advised to compare activations at F3 channel locations between 3-month and 12-month-olds, as F3 may not sample activities from the left inferior frontal gyrus in 3-month-olds.

### Limitation

The current study did not compute thickness of segmentation layers or scalp-to-cortex distances by ROI parcellations. This means that we cannot pinpoint the particular tissue layer(s) or cortical region(s) that may contribute to the age-related changes in scalp-to-cortex distance as done in Beauchamp et al. (2011)’s study. Furthermore, existing evidence indicated that DOT sensitivity as a function of source-detector separation distance was different in GM, WM, CSF, and extracerebral tissues (scalp and skull) in infants and adults (Fukui, Ajichi, & Okada, 2003; Mansouri et al., 2010; Strangman et al., 2013; Strangman, Zhang, & Li, 2014). Future analyses that compare distances from the scalp by tissue types and ROIs could more precisely inform age-related differences in the optimal separation distances for sampling cortical signal changes at different brain regions.

### Conclusions

The current study examined scalp-to-cortex distances and scalp-to-cortical ROI correspondence in infants, children, and adults. There were differences in the scalp-to-cortex distance in infants, which become magnified in children and adults. We found systematic differences in scalp-location-to-anatomical ROI correspondence for different ages, both for spatial projection and for DOT fluence sensitivity. Our findings imply that accurate anatomical interpretations of NIRS/fNIRS data are dependent on developmentally sensitive estimations of DOT sensitivity that account for the head and cortical development. The present study demonstrated that age-appropriate realistic head models can be used with spatial scalp projection and photon migration simulations to provide anatomical guidance for standalone DOT data.

## Acknowledgement

This research was supported by grants from the National Institute of Child Health and Human Development (NICHD-R01-HD18942, NICHD-R37-HD18942, NIHCD-R03-HD091464) to J.E. Richards.

## Code, Data, and Materials Availability

The age-specific average templates, including T1, DOT sensitivity average template can be accessed from the “Neurodevelopmental MRI Database”. The Database is available online: http://jerlab.psych.sc.edu/NeurodevelopmentalMRIDatabase/ for information and https://www.nitrc.org/projects/neurodevdata/ for access. Details on the online access are provided in Richards, Sanchez, Phillips-Meek, and Xie (2015). The codes that support the findings of this study are available from the corresponding author, XF, upon request.

