## Supplemental Information for "Investigating Developmental Changes in Scalp-Cortex Correspondence Using Diffuse Optical Tomography Sensitivity in Infancy"

**Method**

**MRI Segmentation**

Each head MRI volume was segmented into 9 or 10 media types: gray matter (GM), white matter (WM), cerebrospinal fluid (CSF), non-myelinated axons (NMA), other brain matter, skin, skull, air, eyes, and other inside skull material. The FSL FAST procedure (Zhang, Brady, & Smith, 2001) was used to segment the T1-weighted images into GM, WM, or other matter (OM). The GM regions was further separated into gray matter and non-myelinated axons for infants 12 months of age or younger by using the pattern of GM/WM in the average MRI template from two-year-olds as a probability map and identifying participant GM as probable GM or NMA. The CSF was identified in the T2W images using a threshold procedure. The CSF was removed from the materials from the FAST procedure, with the remainder defined as GM, WM, NMA or other inside skull material. The BETSURF procedure (Jenkinson, Pechaud, & Smith, 2005; Smith et al., 2004) was used with the extracted brain, T1W and T2W volumes, to identify skull and scalp regions. The nasal cavity and eyes were identified manually using MRIcron (Rorden, 2012; Rorden & Brett, 2000). Finally, any other matter inside the head volume not defined as above was defined as “other inside skull material”. This generally was in the region of the neck and consisted primarily of muscle and secondarily of spinal bone.

**Mesh Generation**

We generated both sparse and dense “segmented FEM mesh” (also see Method in the main text). The mesh is required only for the MMC package (Fang, 2010). The sparse mesh was used for the MMC when the program did not converge with the dense mesh. The dense “segmented FEM mesh” was used for MCX (Fang & Boas, 2009) and tMCimg (Boas, Culver, Stott, & Dunn, 2002) to find a segment element that is closest to an electrode position. Figure 1 shows the mean numbers of nodes and elements for sparse and dense meshes across age groups. The average number of nodes was 301263, 417523, 440384, for the infants, children, and adults, respectively; average number of elements was 1,752,221, 2,449,375, and 2,582,646; and average tetra volumes were 21, 22, and 27 cubic mm. The change in node and element size reflect increases in head size over these ages.

**10-10 Electrodes Placement**

The 81 10-10 electrode locations were constructed based on the “unambiguously illustrated 10-10 system (Jurcak, Tsuzuki, & Dan, 2007). We divided the electrode positions to six groups for visualization purposes. The Cz was located at the intersection of the front-to-back central curve (Nz to Iz) and the left-to-right central curve (LPA to RPA). The “z” electrodes were placed at the 10% intervals on the Nz-Cz-Iz central curve (group 1). The LPA-RPA central curve was divided in 10% increments to set electrodes T7 to T8. From Nz to LPA to Iz, the N1, I1, and the “9” electrodes (e.g. AF9 and PO9) were identified in 10% intervals (group 2), and likewise for the N2, I2, and the “10” electrodes (e.g. AF10 and PO10) set on the right hemisphere (group 3). The curve Fpz to T7 to Oz was divided in 10% increments to generate Fp1, O1, and the “7” electrodes (group 4), and the same method was applied to identify Fp2, O2, and the “8” electrodes on the right hemisphere (group 5). Lastly, the “z”, “7” and “8” locations were used to define “1” and “2”, ‘3” and “4”, and “5 and “6” electrode locations (group 6).

**Virtual Channel Construction**

We constructed 130 S-D channels from neighboring 10-10 electrode locations based on the fNIRS Optodes’ Location Decider (fOLD) toolbox (Zimeo Morais, Balardin, & Sato, 2018). These channels were centered at the 81 10-10 electrode locations and located at single surrounding 10-10 electrode locations. The fOLD channels were formed for the individual MRIs and age-matched average templates of 3-month-olds (*M*_separation_=23.08mm, *SD*=1.79), 6-month-olds (*M*_separation_=24.41mm, *SD*=1.01), and 20-24-year-olds (*M*_separation_=29.92mm, *SD*=1.59). Figure 2 displays the fOLD channels on the 6-month average template.

**Monte Carlo (MC) Photon Migration Simulations**

The inputs of Monte Carlo simulations include a segmented head model that defines the tissue types. Five tissue types are typically identified: the scalp, skull, cerebrospinal fluid (CSF), gray matter, and white matter (Fukui, Ajichi, & Okada, 2003; Mansouri, L'Huillier, Kashou, & Humeau, 2010; Strangman, Li, & Zhang, 2013). We launched 10^8^ photons for the time window of 0 to 5 nanoseconds. The fluence resolution was 50 time-gates (Zimeo Morais et al., 2018). The wavelength was set at 690 nanometers. Photons were sent from the 358 10-5 electrode positions. The optical properties of the tissue types that are specified include the absorption coefficient, the scattering coefficient, the anisotropy coefficient, and the index of refraction. It should be noted that there has been no consensus on the optical properties in the literature (Zimeo Morais et al., 2018). The values differ considerably across studies (Boas et al., 2002; Brigadoi & Cooper, 2015; Custo et al., 2010; Strangman, Franceschini, & Boas, 2003; Strangman et al., 2013; Whiteman, Santosa, Chen, Perlman, & Huppert, 2017; Zimeo Morais et al., 2018). Different values may be needed for adult and infant-child participants (cf Brigadoi & Cooper, 2015; Fukui et al., 2003; Whiteman et al., 2017). Table 2 summarizes optical properties used in a number of adult and infant-child studies. Our input values for the optical properties of the head media were shown in the Main Text Table 2. They were based on values used in Perdue, Fang, & Diamond (2012), which primarily came from Strangman, Franceschini, & Boas (2003) and Yaroslavsky et al. (2002). These values were also used in Fang (2010).

We used the MC eXtreme (MCX; Fang & Boas, 2009), tMCimg (Boas DA, 2002), and a mesh-based MC method (MMC; Fang, 2010) for modeling photon propagation. The principal distinctions between tMCimg, MCX, and MMC are computational efficiency and the modeling of boundary structure. MCX improves from tMCimg by employing parallel computing in graphics processing units (GPUs) to simultaneously modeling the propagation of a large number of photons. tMCing and MCX are both voxel-based simulations. This means that they discretize boundaries into voxelated, grid-like geometry. This could introduce estimation error of reflection and transmission when a photon reaches certain boundaries. MMC mitigates the problem by using mesh-based head models with tetrahedral elements and ray-tracing calculations. It better approximates complex boundary structures and thus could provide more accurate fluence estimation (Fang, 2010; Tran & Jacques, 2020; Yan, Tran, & Fang, 2019). The input parameters for tMCimg and MMC were the same as those for MCX (see the “Photon Migration Simulations” in Method). MMC was only performed 3-month and 6-month individual MRIs and age-matched average templates.

**DOT Sensitivity Analyses from tMCimg and MMC Outputs**

We computed the “Direct DOT” and “S-D Channel DOT” as described in the “DOT Sensitivity Analyses” section in the Main Text. The Direct DOT distance was defined as the distance from the 10-10 scalp electrode location to the brain voxel with the maximum fluence value. The S-D Channel distance was measured as the distance from the 10-10 channel location (see “Virtual Channel Construction” in the Main Text) to the voxel with the maximum S-D Channel DOT value.

**DOT Sensitivity Analyses for fOLD Channels**

S-D Channel DOT was calculated for each fOLD channel by taking the product of source fluence distribution and the detector fluence distribution. We did not normalize S-D Channel DOT outputs by the sum of fluence for all voxels as described in Zimeo Morais et al. (2018). We also computed the S-D Channel distance for fOLD channels.

**Results**

**Scalp-location-to-ROI Mapping**

Table 3 provides a look-up procedure that maps the scalp electrode and channel locations to the ROIs from the lobar (Fillmore, Richards, Phillips-Meek, Cryer, & Stevens, 2015), Hammer (Heckemann, Hajnal, Aljabar, Rueckert, & Hammers, 2006), and LPBA40 atlas (Shattuck et al., 2008). The spatial scalp projection was used to locate the atlas ROI(s) that intersected with the spherical mask created around the 10-10 electrode location. We also identified ROI(s) in the spherical mask that intersected with the S-D channel DOT fluence distribution (Main Text Figure 3). The percentage of voxels in the ROI is displayed in parentheses. ROIs with less than 10% voxels were not shown.

Table 4 provides a comparable look-up procedure as Zimeo Morais (2018). The table displays the anatomical specificity of a given fOLD channel to the ROI(s). We divided the S-D channel DOT for an ROI by the total channel DOT for all ROIs to compute the specificity (%) to the ROI of a given channel (Zimeo Morais et al., 2018). We additional examined fOLD channel specificity to ROIs from the Brainnetome Atlas (Fan et al., 2016). The atlas provides a microanatomical parcellation of 210 cortical and 36 subcortical subregions. The table displays ROIs with channel specificity greater than or equal to 10%. Channel specificities for the Brainnetome ROIs were smaller since the atlas has more fine-grained parcellations. We listed the Brainnetome ROIs with the largest specificity for a given channel if none of the ROI exceeded the 10% specificity threshold.

**Scalp-to-cortex Distance**

We compared S-D Channel DOT distance computed from the average MRI templates and participant-based individual MRIs. Figure 3A shows that the average distances across channel positions were comparable between head models for each group (infants and toddlers: 2 weeks to 2 years; children: 4 years and 12 years; and adults: 20-24 years). Figure 3B displays the mean S-D Channel DOT distance computed from the average MRI templates by age groups. The distances were similar with those calculated from the individual MRIs. The distances increased from infancy to childhood and to adulthood. There were no systematic age-group differences among infants (cf Main Text Figure 4).

Figure 4 presents the effect of sex on the age-related differences in the three methods of scalp-to-cortex distance estimation. Two-way ANOVAs with age group (infants and toddlers, children, and adults) and sex (male, female) were conducted for each estimation method separately. The findings confirmed the main effect of age in the spatial scalp projection, *F*(2,974)=510.69, *p*<.001, Direction DOT, *F*(2,1051)=1718.07, *p*<.001, and S-D Channel DOT distance, *F*(2,1051)=1255.49, *p*<.001. The main effect of sex was marginally significant for the Direct DOT, *F*(1,1051)=3.47, *p*=.06, and S-D Channel DOT distance, *F*(1,1051)=3.18, *p*=.07. Sex did not moderate the age difference for the Direct DOT, *p*=.17, and S-D Channel DOT distance, *p*=.17. However, there was a significant main effect of sex for the scalp projection measure, *F*(1,974)=7.77, *p*=.005. The age-by-sex interaction was also significant, *F*(2,974)=34.63, *p*=.001. Post-hoc analysis indicated that the sex-difference was significantly for the adult group, *F*(1,974)=15.53, *p*<.001. The distance from the scalp electrode to the cortical surface was greater in males, *M*_males_=13.65, *SD*=2.67, than females, *M*_females_=11.78, *SD*=2.27.

Figure 5 showed a comparison between MCX versus tMCimg estimations of the Direct DOT and Channel DOT. The two methods for simulating photon migrations through the head media produced comparable results.

We compared S-D channel DOT distance across the 10-10 channel locations among estimation methods (MCX, MMC, and tMCimg) and head model types (individual MRIs and age-matched average templates) in the 3-month and 6-month groups (Figure 6). The three estimation methods produced comparable distance estimates across channel locations for 3-month and 6-month individual MRIs. The method-difference was more visible for the average templates than the individual MRIs. There were also head-model-difference across the three estimation methods in a number of channel positions.

Lastly, we compared S-D Channel DOT distance estimated using the fOLD channels with the distance estimated using the 10-10 channels (MCX, MMC, and tMCimg) in 3-month and 6-month individual head models and age-matched average templates (Figure 7). The S-D Channel DOT distance estimates were comparable across methods in 3-month individual head models. However, the mean distance across the fOLD channel was larger than the mean distance across the 10-10 channels in the 6-month individual MRIs. The mean distances differed between individual head models and average templates across estimation methods. The mean distance across fOLD channels in the average templates were higher than the distance across the 10-10 channels in both the average templates and individual head models.

A.

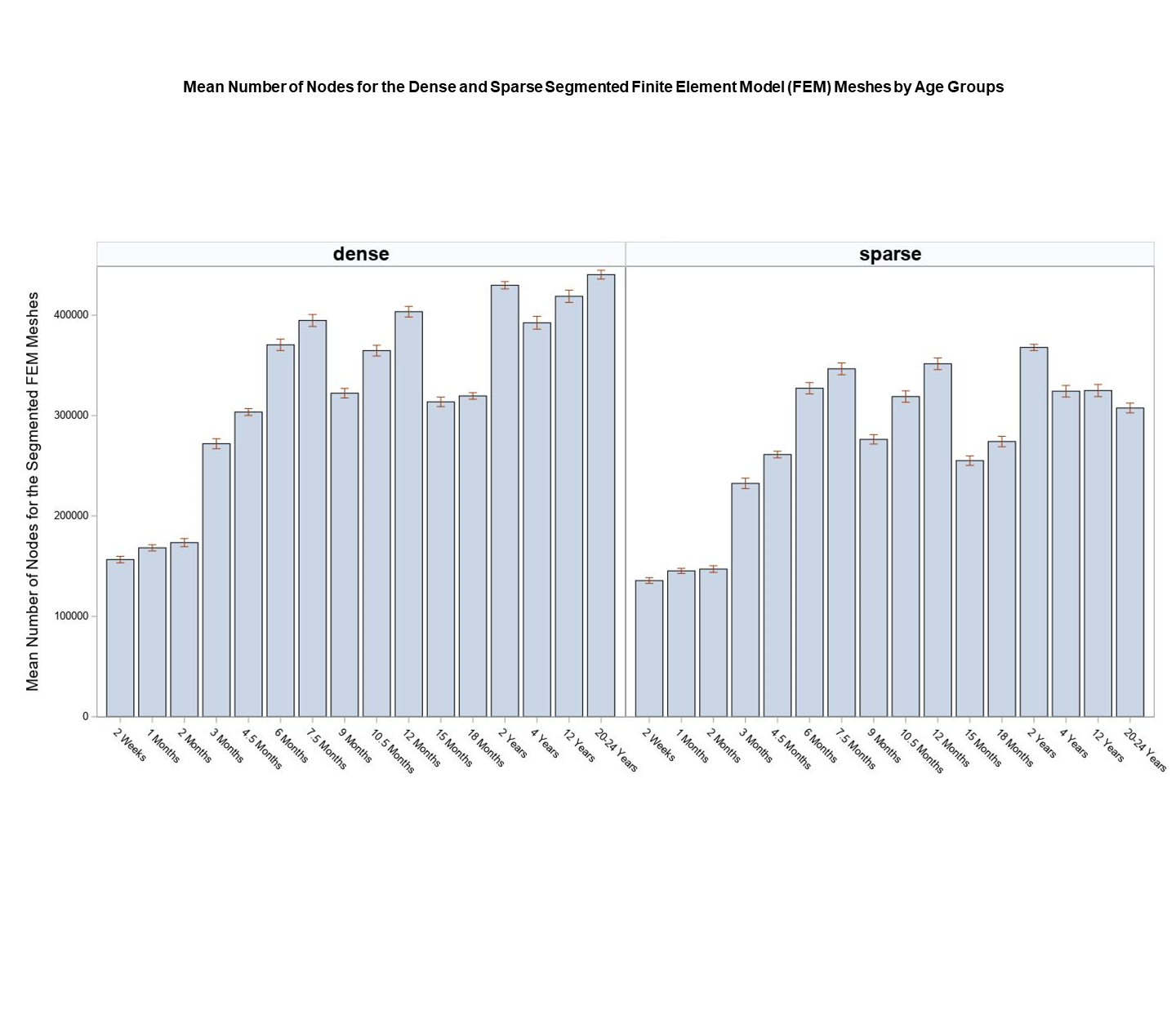

B.

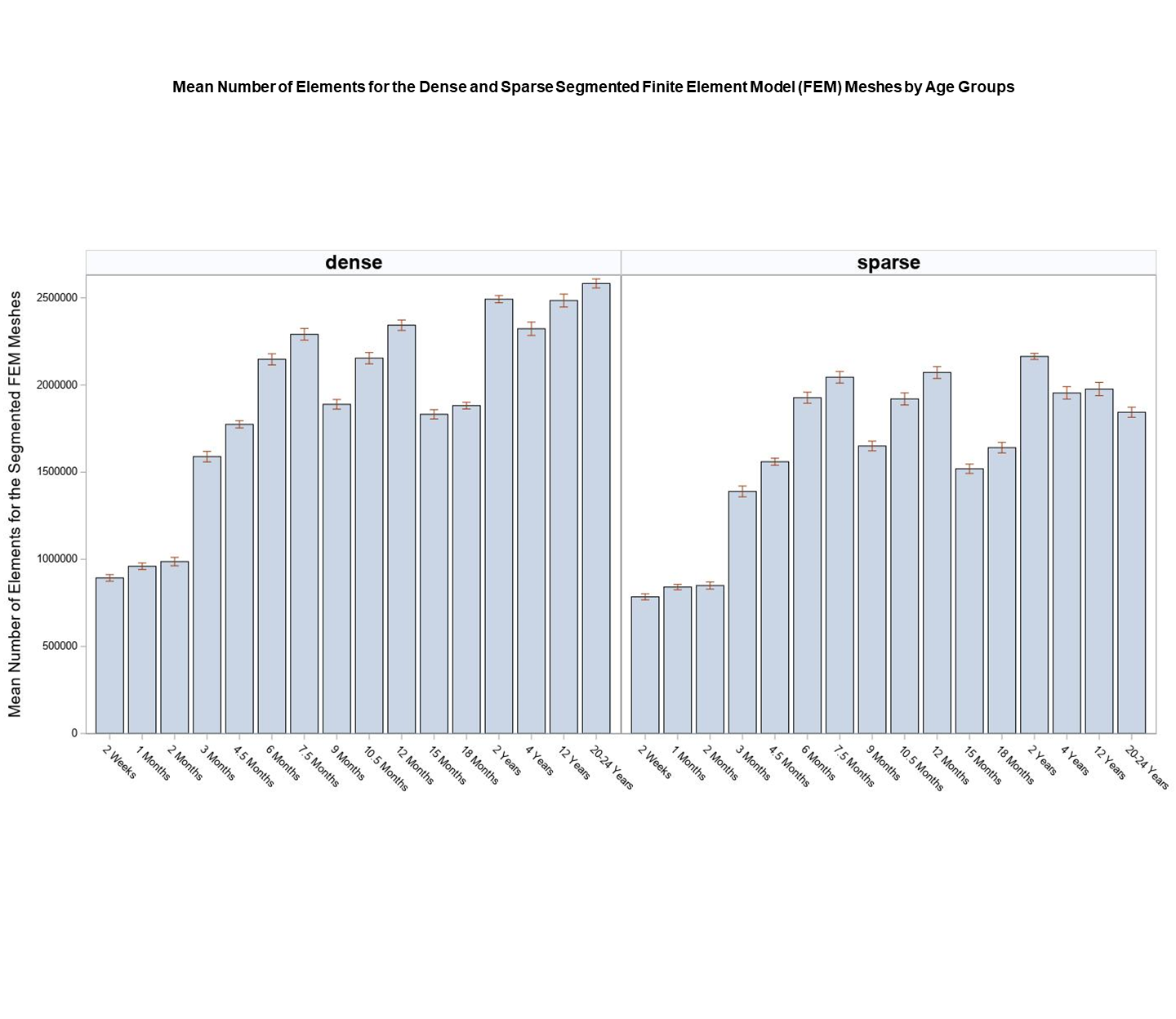

**Fig** 1. Mean number of nodes and elements for the dense and sparse finite element model (FEM) mesh by age groups.

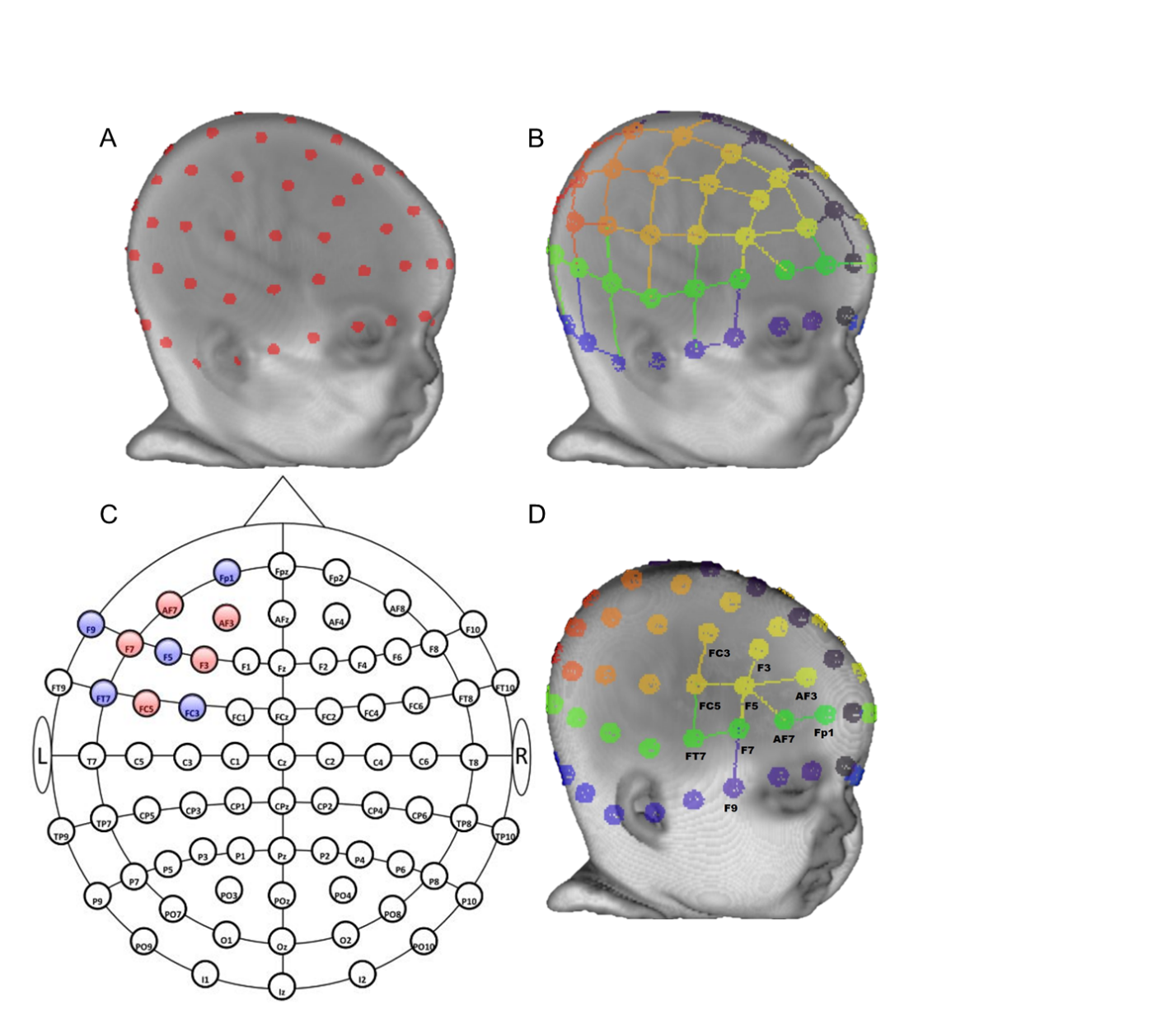

**Fig** 2. Virtual channel placement for fOLD channels. Channels constructed based on the fNIRS Optodes’ Location Decider (fOLD) were displayed on an average template for 6-month infants. A. Ten-ten virtual electrode placement. B. fOLD channel topographical layout. C. Two-dimensional layout of the 10-10 system from the fOLD graphical user interface. It shows the source (red) and detector (blue) combination for the left inferior frontal gyrus from the LONI Probabilistic Brain Atlas (LPBA40; Shattuck et al., 2008). The specificity threshold was set at 1%. D. The topographical layout for channels that are sensitive to the left inferior frontal gyrus with specificity greater than 1%.

A.

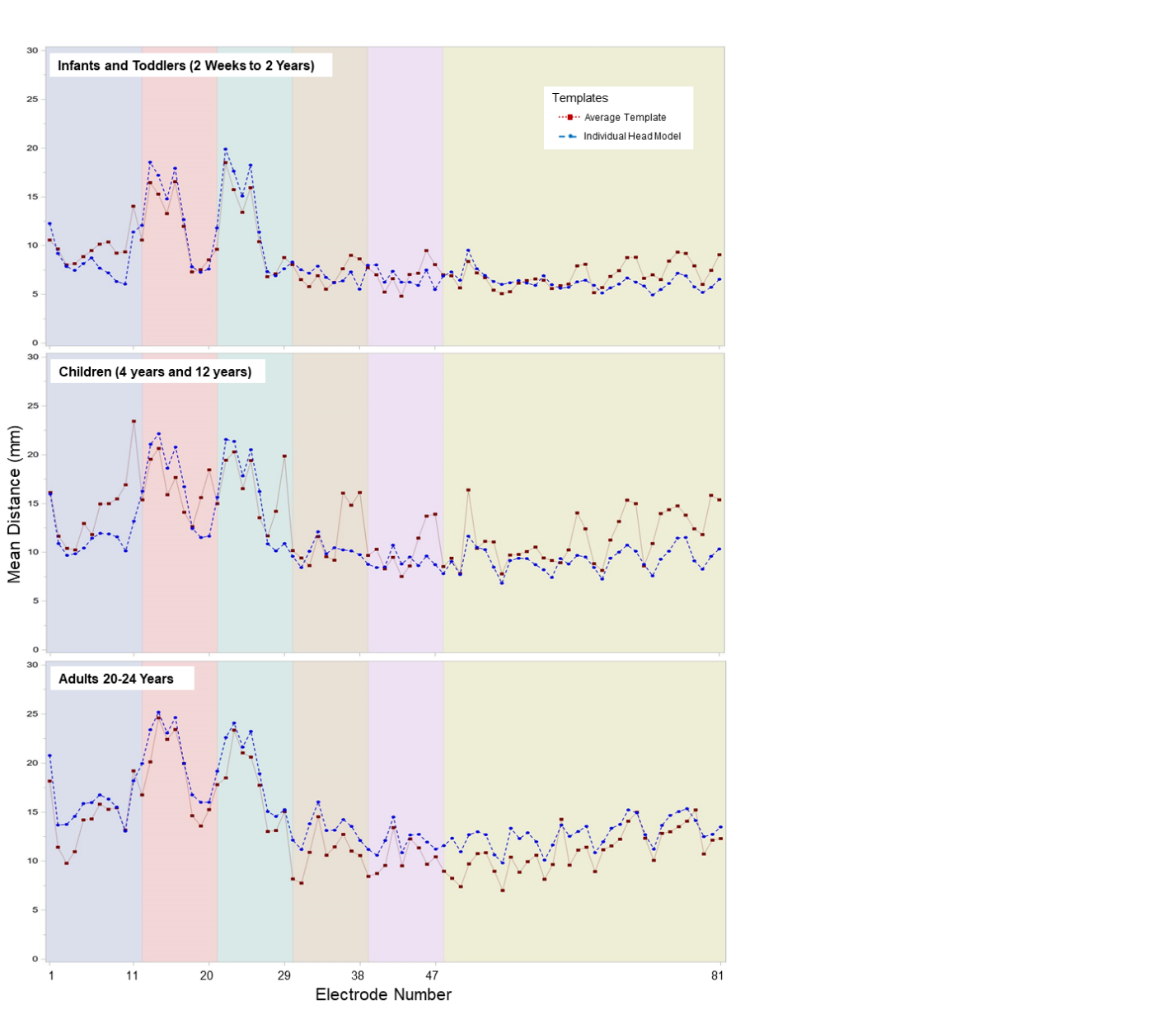

B.

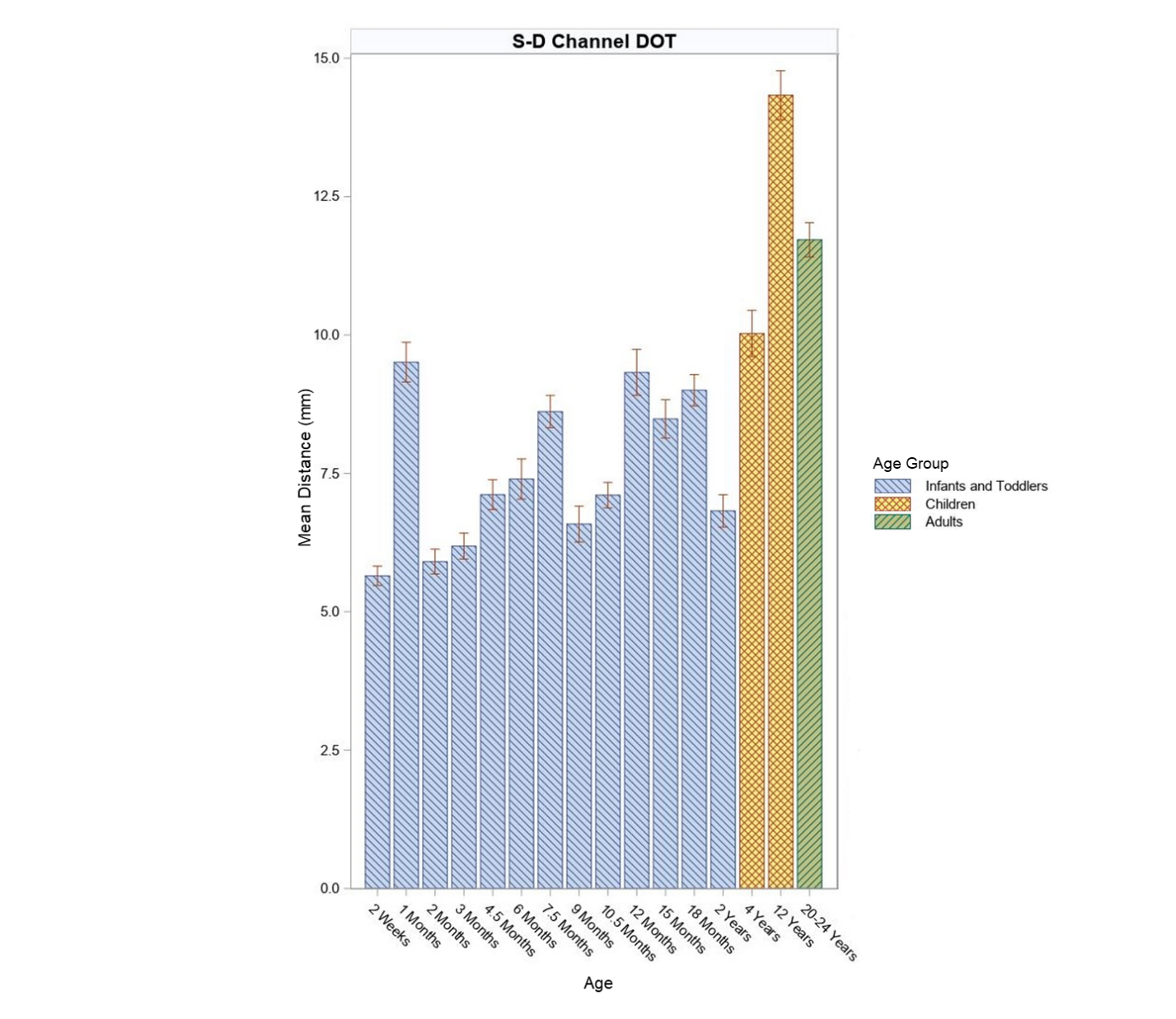

**Fig** 3. Mean Source-Detector (S-D) Channel DOT distances by age groups. A. Mean S-D Channel DOT distances across 10-10 channel locations calculated from age-matched average MRI templates and individual MRIs. B. Mean S-D Channel DOT distances estimated with age-matched average templates by age groups. Data for electrode N1, AF9, F9, FT9, T9/LPA, TP9, N2, AF10, F10, FT10, T10/RPA, and TP10 were excluded.

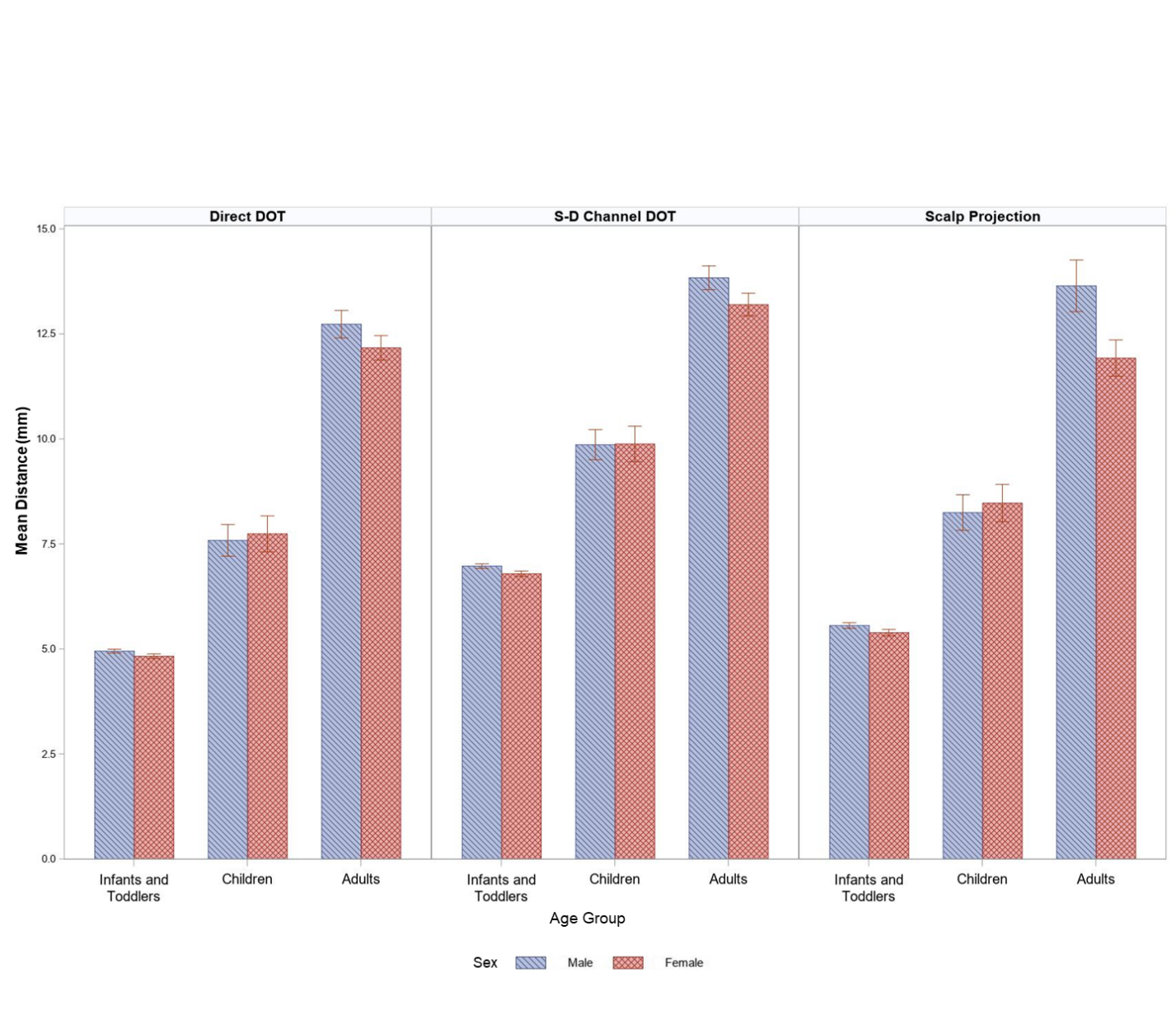

**Fig** 4. Mean scalp-to-cortex distances by sex and age group. Data for electrode N1, AF9, F9, FT9, T9/LPA, TP9, N2, AF10, F10, FT10, T10/RPA, and TP10 were excluded. Direct DOT and S-D Channel DOT were estimated using outputs from the MCX simulations. The infant and toddler group contained age groups from 2 weeks to 2 years. The child group was aged 2 years and 4 years. The adult group was aged 20-24 years.

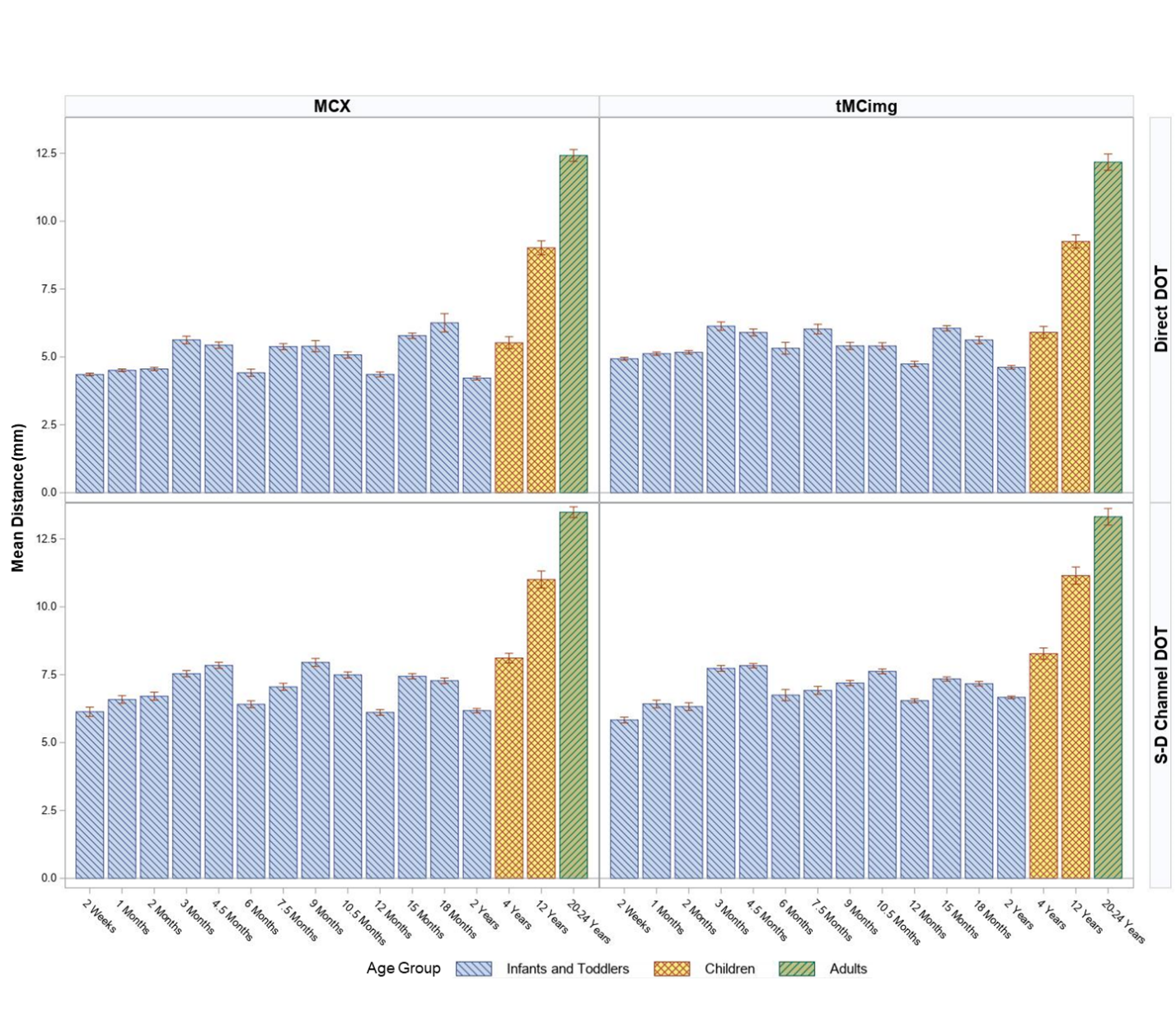

**Fig** 5. Comparison of Direct DOT and S-D Channel DOT distance measures from MCX and tMCimg outputs. Data from electrode and channel position N1, AF9, F9, FT9, T9/LPA, TP9, N2, AF10, F10, FT10, T10/RPA, and TP10 were excluded).

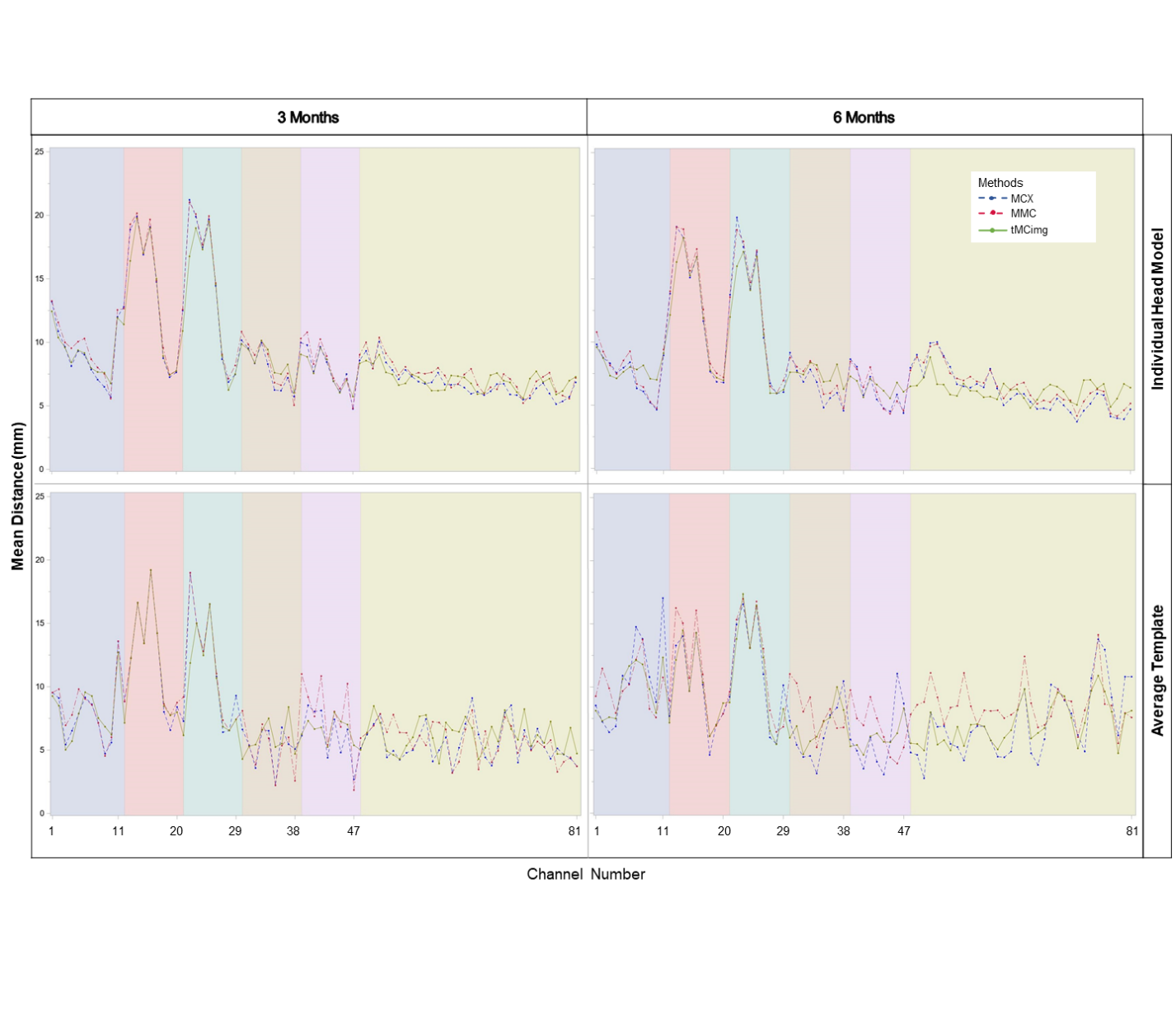

**Fig.** 6. A comparison of S-D channel DOT distance among estimation methods (MCX, MMC, tMCimg) and head model types (individual head models and age-matched average templates) in 3- and 6-month-old infants. Mean distances across 10-10 channel locations are displayed.

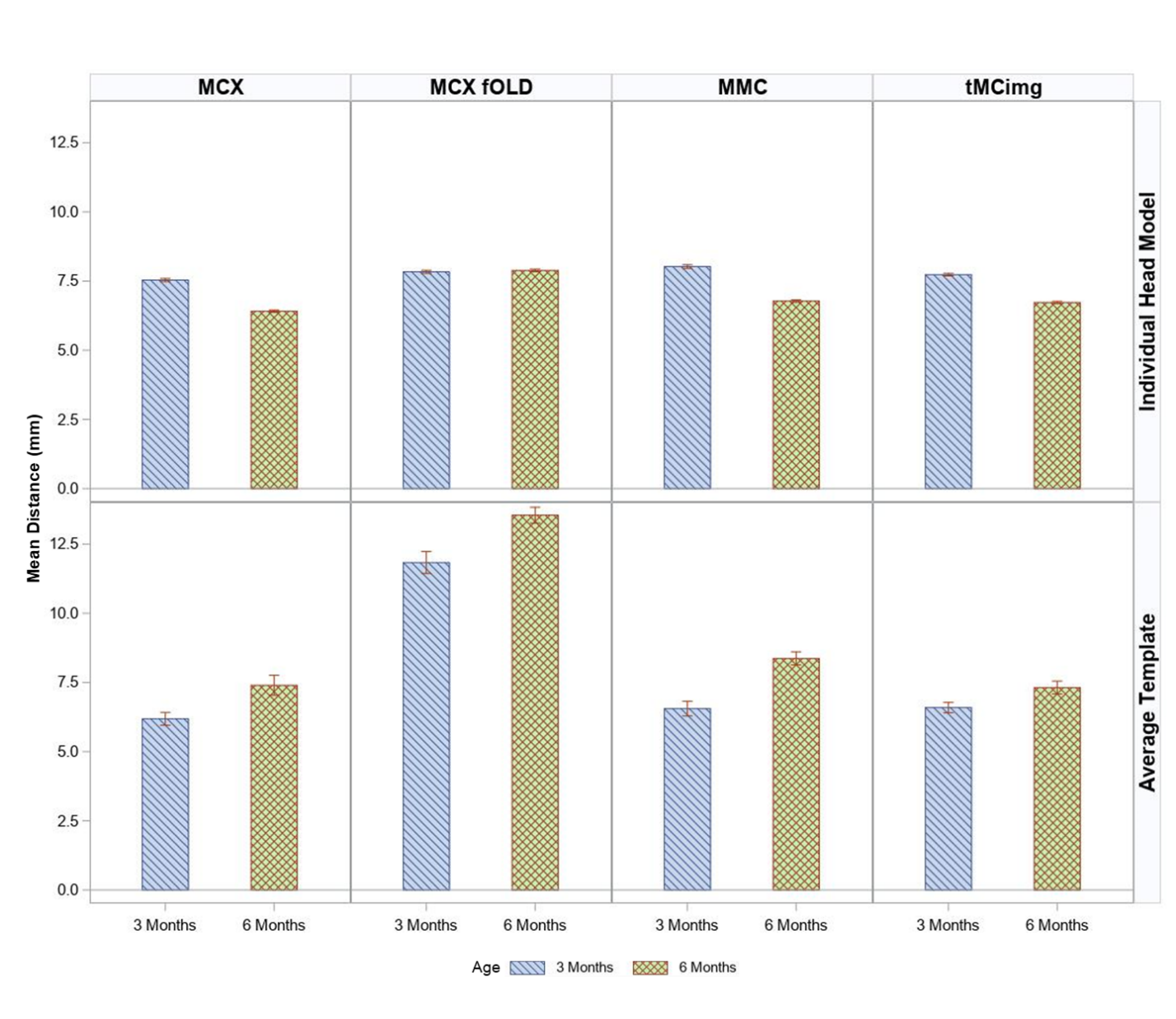

**Fig.** 7. A comparison of S-D channel DOT distance among estimation methods (MCX, MCX with fOLD channels, MMC, tMCimg) and head model types (individual head models and age-matched average templates) in 3- and 6-month-old infants.

**References^[1]^**

Boas DA, C. J., Stott JJ, and Dunn AK. (2002). Three dimensional Monte Carlo code for photon migration through complex heterogeneous media including the adult human head. *Opt Express, 10*, 159-170.

Boas, D. A., Culver, J. P., Stott, J. J., & Dunn, A. K. (2002). Three dimensional Monte Carlo code for photon migration through complex heterogeneous media including the adult human head. *Opt Express, 10*, 159-170.

**Brigadoi, S., & Cooper, R. J. (2015). How short is short? Optimum source-detector distance for short-separation channels in functional near-infrared spectroscopy. *Neurophotonics, 2*(2), 025005. doi:10.1117/1.NPh.2.2.025005**

**Custo, A., Boas, D. A., Tsuzuki, D., Dan, I., Mesquita, R., Fischl, B., . . . Wells, W., 3rd. (2010). Anatomical atlas-guided diffuse optical tomography of brain activation. *Neuroimage, 49*(1), 561-567. doi:10.1016/j.neuroimage.2009.07.033**

**Dehaes, M., Kazemi, K., Pélégrini-Issac, M., Grebe, R., Benali, H., & Wallois, F. (2013). Quantitative effect of the neonatal fontanel on synthetic near infrared spectroscopy measurements. *Human Brain Mapping, 34*(4), 878-889. doi:10.1002/hbm.21483**

Fan, L., Li, H., Zhuo, J., Zhang, Y., Wang, J., Chen, L., . . . Jiang, T. (2016). The Human Brainnetome Atlas: A New Brain Atlas Based on Connectional Architecture. *Cerebral Cortex, 26*(8), 3508-3526. doi:10.1093/cercor/bhw157

**Fang, Q. (2010). Mesh-based Monte Carlo method using fast ray-tracing in Plücker coordinates. *Biomedical Optics Express, 1*(1), 165-175. doi:10.1364/boe.1.000165**

Fang, Q., & Boas, D. A. (2009). Monte Carlo simulation of photon migration in 3D turbid media accelerated by graphics processing units. *Opt Express, 17*(22), 20178-20190. doi:10.1364/oe.17.020178

Fillmore, P. T., Richards, J. E., Phillips-Meek, M. C., Cryer, A., & Stevens, M. (2015). Stereotaxic Magnetic Resonance Imaging Brain Atlases for Infants from 3 to 12 Months. *Developmental Neuroscience, 37*(6), 515-532. doi:10.1159/000438749

**Fukui, Y., Ajichi, Y., & Okada, E. (2003). Monte Carlo prediction of near-infrared light propagation in realistic adult and neonatal head models. *Applied Optics, 42*(16), 6. Retrieved from** [**https://doi.org/10.1364/AO.42.002881**](https://doi.org/10.1364/AO.42.002881)

Heckemann, R. A., Hajnal, J. V., Aljabar, P., Rueckert, D., & Hammers, A. (2006). Automatic anatomical brain MRI segmentation combining label propagation and decision fusion. *Neuroimage, 33*(1), 115-126. doi:<https://doi.org/10.1016/j.neuroimage.2006.05.061>

**Huppert, T. J., Diamond, S. G., Franceschini, M. A., & Boas, D. A. (2009). HomER: a review of time-series analysis methods for near-infrared spectroscopy of the brain. *Applied Optics, 48*, D280-D298.**

Jenkinson, M., Pechaud, M., & Smith, S. M. (2005). BET2: MR-based estimation of brain, skull and scalp surfaces. *In Eleventh Annual Meeting of the Organization for Human Brain Mapping*.

Jurcak, V., Tsuzuki, D., & Dan, I. (2007). 10/20, 10/10, and 10/5 systems revisited: their validity as relative head-surface-based positioning systems. *Neuroimage, 34*(4), 1600-1611. doi:10.1016/j.neuroimage.2006.09.024

Mansouri, C., L'Huillier, J.-p., Kashou, N. H., & Humeau, A. (2010). Depth sensitivity analysis of functional near-infrared spectroscopy measurement using three-dimensional Monte Carlo modeling-based magnetic resonance imaging. *Lasers in Medical Science, 25*(3), 431-438. doi:<http://dx.doi.org/10.1007/s10103-010-0754-4>

**Perdue, K. L., Fang, Q., & Diamond, S. G. (2012). Quantitative assessment of diffuse optical tomography sensitivity to the cerebral cortex using a whole-head probe. *Physics in medicine and biology, 57*(10), 2857. doi:10.1088/0031-9155/57/10/2857**

Rorden, C. (2012). MRIcroGL. *Retrieved from McCausland Center:* [*http://www.mccauslandcenter.sc.edu/mricrogl/*](http://www.mccauslandcenter.sc.edu/mricrogl/).

Rorden, C., & Brett, M. (2000). Stereotaxic Display of Brain Lesions. *Behavioural Neurology, 12*, 421719. doi:10.1155/2000/421719

Shattuck, D. W., Mirza, M., Adisetiyo, V., Hojatkashani, C., Salamon, G., Narr, K. L., . . . Toga, A. W. (2008). Construction of a 3D probabilistic atlas of human cortical structures. *Neuroimage, 39*(3), 1064-1080. doi:<https://doi.org/10.1016/j.neuroimage.2007.09.031>

Smith, S. M., Jenkinson, M., Woolrich, M. W., Beckmann, C. F., Behrens, T. E. J., Johansen-Berg, H., . . . Matthews, P. M. (2004). Advances in functional and structural MR image analysis and implementation as FSL. *Neuroimage, 23*, S208-S219. doi:<https://doi.org/10.1016/j.neuroimage.2004.07.051>

**Strangman, G., Franceschini, M. A., & Boas, D. A. (2003). Factors affecting the accuracy of near-infrared spectroscopy concentration calculations for focal changes in oxygenation parameters. *Neuroimage, 18*(4), 865-879. doi:10.1016/s1053-8119(03)00021-1**

Strangman, G., Li, Z., & Zhang, Q. (2013). Depth Sensitivity and Source-Detector Separations for Near Infrared Spectroscopy Based on the Colin27 Brain Template. *PLOS ONE, 8*(8), e66319. doi:10.1371/journal.pone.0066319

**Strangman, G., Zhang, Q., & Li, Z. (2014). Scalp and skull influence on near infrared photon propagation in the Colin27 brain template. *Neuroimage, 85*, 136-149. doi:**[**https://doi.org/10.1016/j.neuroimage.2013.04.090**](https://doi.org/10.1016/j.neuroimage.2013.04.090)

Tran, A. P., & Jacques, S. (2020). Modeling voxel-based Monte Carlo light transport with curved and oblique boundary surfaces. *Journal of Biomedical Optics, 25*(2), 025001. Retrieved from <https://doi.org/10.1117/1.JBO.25.2.025001>

**Whiteman, A., Santosa, H., Chen, D., Perlman, S., & Huppert, T. (2017). Investigation of the sensitivity of functional near-infrared spectroscopy brain imaging to anatomical variations in 5- to 11-year-old children. *Neurophotonics, 5*(1), 011009.** Retrieved from <https://doi.org/10.1117/1.NPh.5.1.011009>

<https://www.ncbi.nlm.nih.gov/pmc/articles/PMC5601503/pdf/NPh-005-011009.pdf>

Yan, S., Tran, A. P., & Fang, Q. (2019). Dual-grid mesh-based Monte Carlo algorithm for efficient photon transport simulations in complex three-dimensional media. *Journal of Biomedical Optics, 24*(2), 020503. Retrieved from <https://doi.org/10.1117/1.JBO.24.2.020503>

**Yaroslavsky, A. N., Schulze, P. C., Yaroslavsky, I. V., Schober, R., Ulrich, F., & Schwarzmaier, H. J. (2002). Optical properties of selected native and coagulated human brain tissues in vitro in the visible and near infrared spectral range. *Phys Med Biol, 47*(12), 2059-2073.** Retrieved from <https://www.ncbi.nlm.nih.gov/pubmed/12118601>

Zhang, Y., Brady, M., & Smith, S. (2001). Segmentation of brain MR images through a hidden Markov random field model and the expectation-maximization algorithm. *IEEE Trans Med Imaging, 20*(1), 45-57. doi:10.1109/42.906424

**Zimeo Morais, G. A., Balardin, J. B., & Sato, J. R. (2018). fNIRS Optodes’ Location Decider (fOLD): a toolbox for probe arrangement guided by brain regions-of-interest. *Scientific Reports, 8*(1), 3341. doi:10.1038/s41598-018-21716-z**

1. References for optical properties summarized in Supplemental Information Table 2 are bolded.
